# Microglial sTREM2 limits dyskinesia and acts on TrkB to support circuit plasticity

**DOI:** 10.64898/2026.05.06.723382

**Authors:** Caitlin Castagnola, Roberta Marongiu, Yuansong Wan, Eileen Ruth S. Torres, Li Fan, Minwoo Wendy Jang, Qiaoling Cui, Jihye Kim, Pearly Ye, Jingjie Zhu, Alisa Panichkina, Alexandra B. Fall, Kendra Norman, Beichen Gao, Nessa Foxe, Man Ying Wong, Daphne Zhu, Maitreyee Bhagwat, Shiaoching Gong, Anna G. Orr, Francis S. Lee, D. James Surmeier, Michael G. Kaplitt, Li Gan

## Abstract

Microglia continuously survey the brain and shape neuronal activity, but their contribution to experience-dependent synaptic plasticity is unclear. Levodopa-induced dyskinesia (LID) is a disabling complication of late-stage Parkinson’s disease (PD) that is linked to maladaptive striatal remodeling and is often assumed to reflect detrimental neuroinflammation. Here we identify a dyskinesia-associated microglial gene program in the striatum of PD patients and show that microglia instead act as a protective brake on LID. In a mouse model, microglial depletion exacerbated dyskinesia, whereas microglial repopulation mitigated it. Delivery of AAV expressing soluble TREM2 (sTREM2) similarly reduced LID without impairing the therapeutic benefit of levodopa. Single-nucleus transcriptomics revealed that microglial loss drives extensive remodeling of both direct and indirect spiny projection neurons (SPNs), while repopulation or sTREM2 reverses a large fraction of LID-associated transcriptional changes. Mechanistically, sTREM2 directly engages TrkB and potentiates BDNF-dependent TrkB-ERK signaling, consistent with positive allosteric modulation. Functionally, sTREM2 enhances BDNF-TrkB-dependent hippocampal synaptic plasticity and acutely rebalances striatal dendritic excitability in a compartment- and cell type-specific manner. These findings reveal an unexpected neuroimmune pathway in which microglia restrain maladaptive plasticity via sTREM2-TrkB signaling, with therapeutic implications.

## Main Text

Microglia provide a critical interface between immune signaling and brain circuit function(*1–3*). Neuroinflammatory states disrupt this dialogue. For example, immune activation enhances seizure vulnerability, impairs inhibitory synapse maintenance, and perturbs synaptic plasticity(*3–6*). In Parkinson’s disease (PD)(*7–9*), inflammation coincides with increased excitability and characteristic oscillatory abnormalities, suggesting that immune-driven microglial changes contribute to network pathophysiology.

One of the most clinically important network pathophysiological states in PD is levodopa-induced dyskinesia (LID). In its early stages, PD is effectively treated with levodopa (L-DOPA), but as the disease progresses and the dose of L-DOPA needed to achieve symptomatic benefit rises, it commonly results in LID, compromising patient’s quality of life (*10,11*). Although the mechanisms underlying LID are multifaceted, maladaptive plasticity in striatal direct and indirect SPNs (dSPNs and iSPNs, respectively), is critical. Although inflammation is associated with LID, its role in the aberrant striatal adaptations causing LID is unclear (*12–17*).

A major microglial effector of network homeostasis is triggering receptor expressed on myeloid cells 2 (TREM2). The extracellular domain of TREM2 is shed to generate soluble TREM2 (sTREM2) (*18,19*). While biofluid sTREM2 levels reflect TREM2 linked neuroinflammatory activity, higher baseline cerebrospinal fluid sTREM2 is associated with slower cognitive decline and reduced conversion from mild cognitive impairment to dementia (*20–22*), supporting protective functions of sTREM2. However, how sTREM2 acts on neurons remains largely unknown.

Here, we identify a microglial gene expression signature in PD patients and observe a similar microglial transcriptional signature in a mouse model with LID. In this model, we show that microglia actively constrain dyskinetic behavior, and that their depletion exacerbates LID. Using AAV-mediated delivery, we find that sTREM2 attenuates LID without diminishing the therapeutic benefit of L-DOPA. Using co-IP and split-luciferase assay, we found that sTREM2 binds to TrkB, and potentiates signaling induced by the endogenous TrkB ligand BDNF and enhances TrkB-dependent synaptic plasticity. Moreover, sTREM2 mimics the effects of TrkB activation on the dendritic activity of SPNs, modulating the striatal circuitry in a manner consistent with its attenuation of LID. These findings establish microglia as positive regulators of striata circuits underlying LID and identify sTREM2-TrkB signaling as an unexpected mediator of this regulation.

### Identification of gene co-expression modules in human PD patients that are preserved across species

To investigate microglial contributions to LID in humans, we performed weighted gene co-expression network analysis on a published bulk RNA sequencing dataset from the caudate and putamen of healthy controls and patients with PD(*23*). From the top 5% highly expressed genes, we identified seven co-expression modules with distinct eigengene structures (Fig.1A and Fig. S1A). Out of these gene modules, onl the black module was significantly correlated with dyskinesia in PD patients (Fig. 1B). This module was enriched for phagocytosis-related and microglia-associated genes (C1QA, C1QB, C1QC, C3, ITGB2, TREM2, TMEM119, CSF1R) (Fig. 1C, D and Fig. S1B). Notably, other modules also contained inflammatory or homeostatic microglia genes, suggesting that multiple microglia programs may contribute to different clinical features of PD (Fig. S1C and Table S1).

**Fig. 1.**
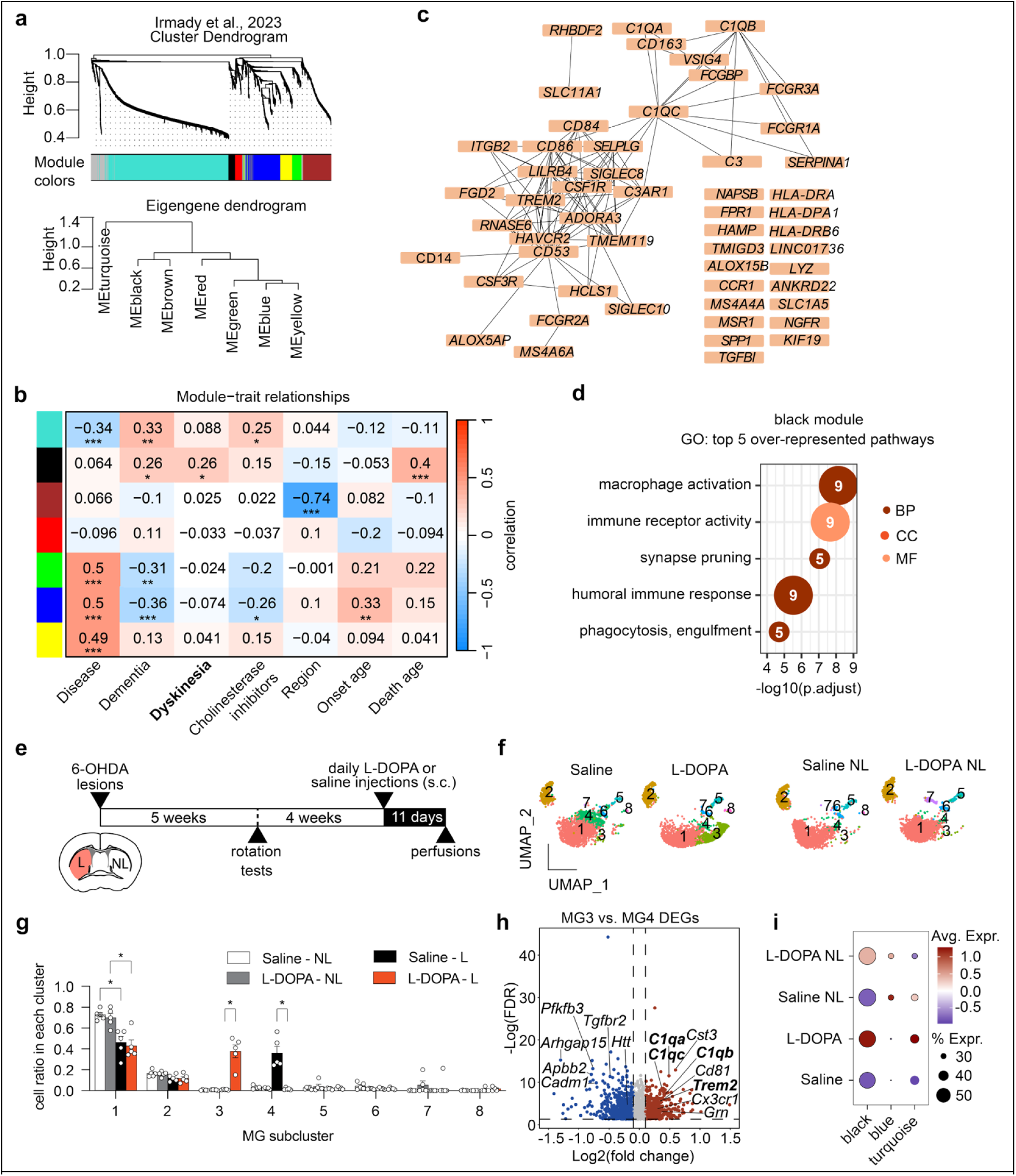
LID alters microglial states in PD patients and mouse models. (**a**) Dendrogram showing the hierarchical clustering of individual genes into their designated co-expression modules. Modules are represented by colors. ME, module eigengene. (**b**) Heatmap of correlation between module eigengenes and traits. Printed values are the Pearson correlation coefficient between the corresponding module and trait. \**p*<0.05, \*\**p*<0.01, \*\*\**p*<0.001, statistics are FDR-adjusted p-values derived from the WGCNA package. (**c**) Network graph showing the strongest correlations between genes in the black module. (**d**) Over-representation analysis of GO terms using the genes in the black module. (**e**) Schematic of the unilateral 6-OHDA mouse model timeline. The striatum from both lesioned (L) and non-lesioned (NL) hemispheres was dissected for snRNA-sequencing. (**f**) UMAP of microglia subclusters split by condition. Colors correspond to microglia subclusters. (**g**) Bar plot of the distribution of microglia (MG) assigned to each subcluster within each condition. Each dot represents the proportion of microglia in that subcluster in one sample (n = 5 striatal samples per group from 10 mice; mean ± s.e.m.). \**p*<0.05, repeated measures 2-way ANOVA with Tukey’s post hoc multiple comparisons test. (**h**) Volcano plot of DEGs between subclusters MG3 versus MG4 with select genes highlighted. Dashed lines indicate FDR thresholds of *p*<0.05 and log2(fold change) thresholds of ±0.1. Red values indicate higher expression in MG3. (**i**) Dot plot of projected module eigengene values in mouse microglia split by condition.

We next asked whether this dyskinesia-linked module was preserved in striatal microglia from the well-established 6-OHDA mouse model with LID. Briefly, this mouse model involves unilateral injection of the 6-OHDA neurotoxin into the medial forebrain bundle to selectively kill dopaminergic nigrostriatal cells and generate hemiparkinsonism (*24*). Following recovery and *in vivo* lesion confirmation with the apomorphine-induced rotation test, daily administration of high concentrations of L-DOPA can induce abnormal involuntary movements characteristic of LID (*24*) (Fig. 1E). Tissues were harvested during the on-state of LID, 1 hour after the last injection of L-DOPA. After post-mortem confirmation of lesioning and microglia density increases (Fig. S2), we performed single nucleus RNA sequencing (snRNA-seq) of lesioned and non-lesioned striata (Fig. 1F, G and Fig. S3). Pseudobulk analysis showed that the LID-associated subcluster (MG3) expressed higher levels of mouse orthologs corresponding to key black module genes, including *C1qa, C1qb, C1qc,* and *Trem2* (Fig. 1H and Table S2). To assess conservation quantitatively, we mapped human genes to mouse orthologs and performed module preservation analysis (*25,26*). Three modules (turquoise, blue, and black) showed moderate preservation in mouse microglia (Fig. S4A). After accounting for module size using median rank statistics, the black module exhibited the strongest preservation and showed the highest relative eigengene values in microglia from the lesioned, L-DOPA-treated striatum (Fig. 1I and Figure S4B-D). These findings implicate microglia in the regulation of dyskinesia across mouse models and human disease.

### Microglia protect against LID and partially reverse SPN transcriptional changes

To determine the role of microglia in LID, we depleted microglia with the CSF1R inhibitor PLX5622 at the start of LID induction and assessed dyskinesia scores weekly (Fig. 2A). Microglia depletion did not alter baseline locomotion but significantly worsened dyskinesia, with elevated dyskinesia scores throughout the first hour after L-DOPA administration across all treatment weeks (Fig. 2B-D). Returning mice to control chow after depletion restored microglia and partially rescued dyskinesia severity (Fig. 2C,D) indicating that microglia presence confers protection against LID.

**Fig. 2.**
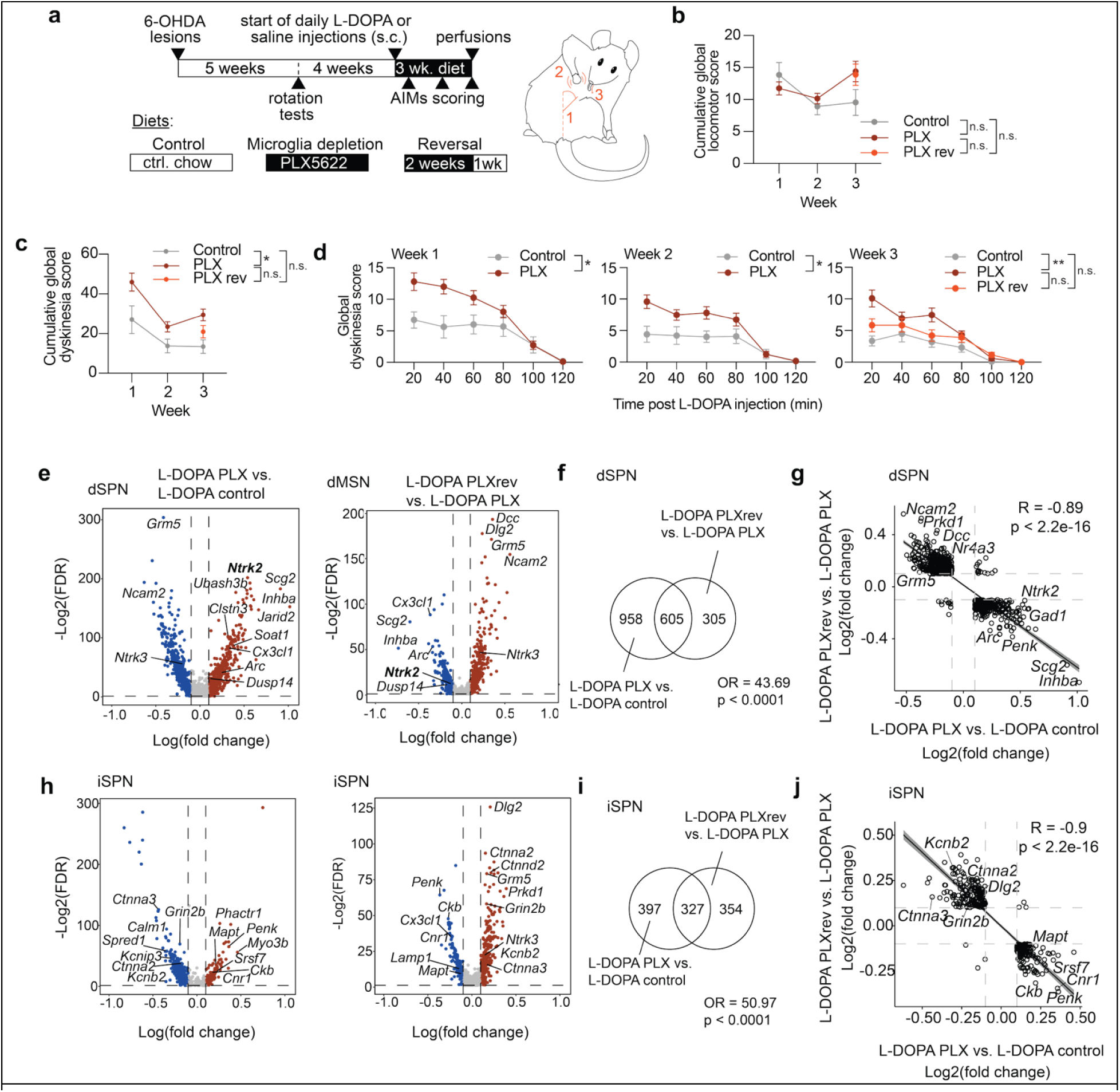
Microglia protect against LID and partially reverse LID-induced SPN transcriptomic changes. (**a**) Schematic of the timeline for the LID model with microglia depletion and diagram of dyskinetic behaviors scored (right). Diet groups are listed (bottom) and the timeline for each is depicted by color of the bar, with control chow in white and PLX-infused chow in black. Dyskinesia scores are made from assessment of axial (1), limb (2), and orolingual (3) dyskinesias. (**b**) Cumulative non-dyskinetic locomotion scores across weeks, groups are denoted by color. (**c**) Cumulative dyskinesia scores across weeks, groups are denoted by color. \*\**p* < 0.05, repeated measures mixed-effects model with Sidak’s post hoc multiple comparisons test. (**d**) Dyskinesia scores measured at scoring intervals after L-DOPA administration within each behavior session for all three weeks. Each dot represents the average (± s.e.m.) score from all mice in the group (n = 11 mice on control diet; n = 25 mice on the PLX diet for weeks 1 and 2, 13 mice on PLX diet for week 3; n = 12 mice on the reversal diet for week 3). \**p* < 0.05, \*\**p* < 0.01, repeated-measures 2-way ANOVA. **(e)** Volcano plots of dSPN DEGs comparing L-DOPA control versus L-DOPA PLX diets (left) and L-DOPA PLX reversal versus L-DOPA PLX diets (right), with select genes highlighted. **(f)** Venn diagrams showing overlapping dSPN DEGs from the two comparisons. **(g)** Scatter plot of shared dSPN DEGs with linear regression and 95 percent confidence interval. **(h)** Volcano plots of iSPN DEGs comparing L-DOPA control versus L-DOPA PLX diets (left) and L-DOPA PLX reversal versus L-DOPA PLX diets (right), with select genes highlighted. **(i)** Venn diagrams showing overlapping iSPN DEGs from the two comparisons. **(j)** Scatter plot of shared iSPN DEGs with linear regression and 95% confidence interval. Significance and odds ratios were determined with the GeneOverlap(v1.38.0) package (**f, i**). Pearson correlation coefficients and p-values were determined with the ggpubr(v0.6.1) package (**g, j**).

LID is temporally linked to the L-DOPA on-state, when peak dopaminergic stimulation drives maladaptive striatal plasticity and dyskinesia. We therefore profiled the lesioned striatum 1 hour after the final L-DOPA injection—capturing the on-state—using snRNA-seq (Fig. S5). Subclustering dSPNs revealed an L-DOPA-associated population (dSPN2) whose markers significantly overlapped and correlated with a published LID dSPN signature (*27*), and included immediate early genes associated with increased neuronal activity (*Arc, Fos, Fosb, Egr1*; Fig. S6A-D). Microglia depletion did not change the proportion of L-DOPA responsive dSPN2 nuclei but induced substantial transcriptional remodeling: pseudobulk analysis identified 1563 differentially expressed genes (DEGs) in dSPNs (Fig. 2E,F and Figure S6A,B). Pseudobulk analysis revealed that many microglia depleted DEGs in dSPNs showed a reversal of expression patterns upon microglia repopulation (Fig. 2E-G). A similar pattern emerged in iSPNs, albeit with a different set of DEGs (Fig. 2H-J). That is, many genes upregulated in SPNs during microglia depletion were downregulated upon repopulation, while genes suppressed during depletion increased during repopulation (Fig. 2G, J).

Among the microglia-sensitive DEGs identified in SPNs were many genes previously implicated in rodent models of LID (*28,29*). Notably, TrkB (*Ntrk2*) was among the most strongly upregulated transcripts, whereas metabotropic glutamate receptor 5 (mGluR5; *Grm5*) was among the most significantly downregulated in dSPNs following microglial depletion (Fig. 2E,G). These two receptors exert opposing effects in LID: TrkB deletion exacerbates dyskinesia, whereas pharmacologic antagonism of mGluR5 confers protection (*30–35*). Together, these findings indicate that microglia—or microglia-derived factors—profoundly modulate key signaling pathways governing SPN imbalance, a hallmark of dyskinesia.

### Intravenous injection of AAV-sTREM2 ameliorates LID without interfering with L-DOPA efficacy

Given that TREM2 is a major microglia receptor whose shed ectodomain (sTREM2) has been linked to neuroprotective functions in neurodegenerative settings, we asked whether increasing systemic sTREM2 could counteract dyskinesia. To raise sTREM2 levels non-invasively, we delivered a brain-permeable AAV expressing sTREM2 (or GFP control) intravenously into 6-OHDA–lesioned mice (Fig. 3A). Postmortem ELISA and immunofluorescence confirmed robust and widespread sTREM2 levels in the cortex (Fig. 3B) and striatum (Fig. 3C).

**Fig. 3.**
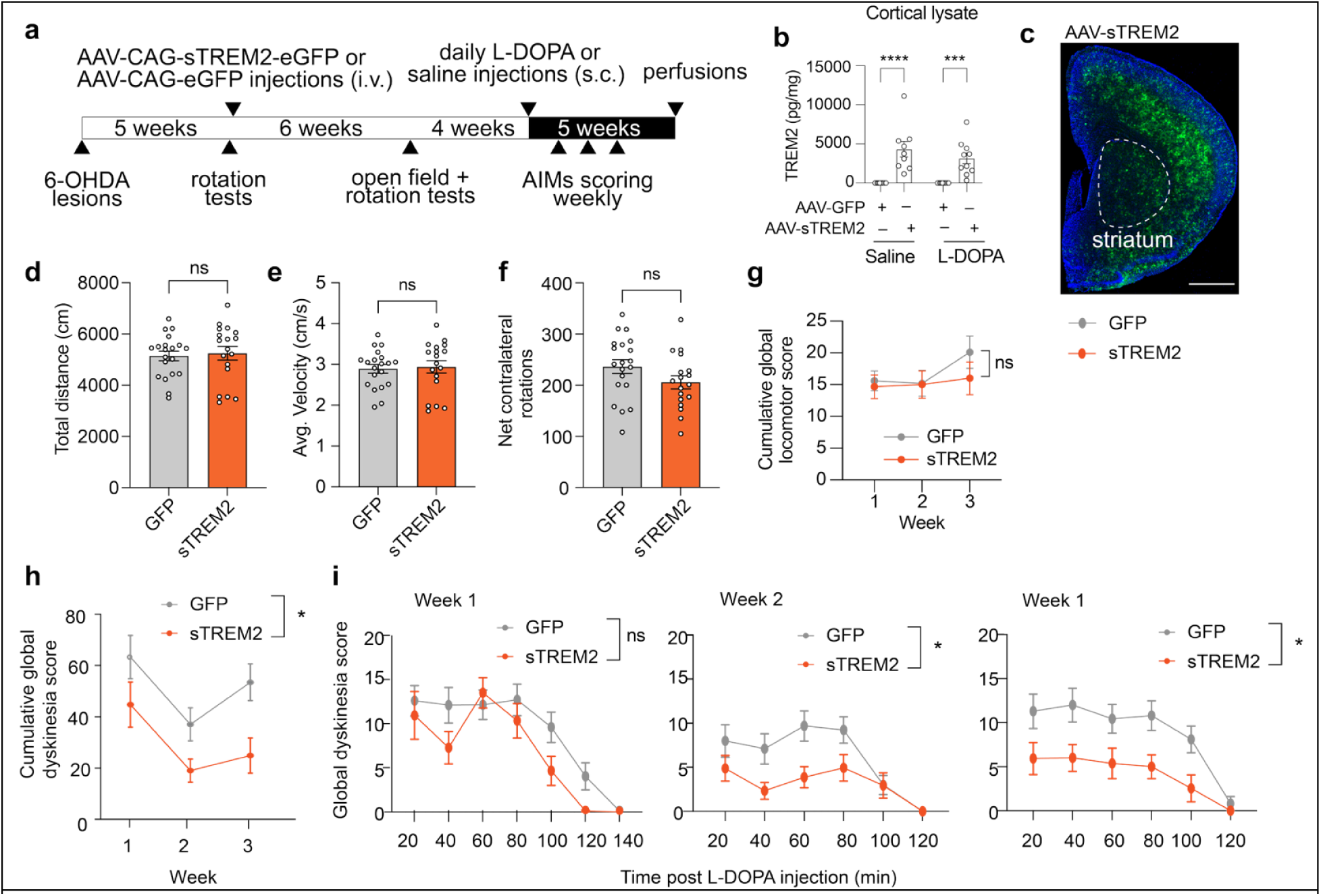
Intravenous injection of AAV-sTREM2 ameliorates LID without affecting general locomotion. **(a)** Timeline for the LID paradigm with AAV-mediated sTREM2 expression. **(b–c)** ELISA confirmation of increased TREM2 protein in cortex and plasma from sTREM2 injected mice compared to GFP controls (**b**) and in striatum using IHC (**c**) (one sTREM2 mouse excluded for insufficient expression; n=10 saline-treated GFP-injected; n=10 L-DOPA-treated GFP-injected; n=10 L-DOPA-treated sTREM2-injected; n=9 saline-treated sTREM2-injected mice). ***p < 0.001, \*\*\*\***p < 0.0001,** two-way ANOVA with Sidak’s post hoc test. Scale bar, 1000μm. **(d–e)** Total distance (**d**) and velocity (**e**) in the open field before LID induction. **(f)** Net contralateral rotations in the apomorphine test before LID induction. ns, not significant; unpaired two-tailed t test; n = 20 GFP-injected and 19 sTREM2-injected mice (**d–f**). **(g)** Cumulative non-dyskinetic locomotion scores during L-DOPA treatment. **(h)** Cumulative global dyskinesia scores. **(i)** Global dyskinesia scores across scoring intervals for weeks 1–3. Data shown as mean ± s.e.m. Each dot represents average cumulative global motor scores from mice in that group (**g–i**). ns, not significant; *p < 0.05 main effect of sTREM2, repeated measures two-way ANOVA (**g–i**).

We first evaluated whether sTREM2 affected baseline motor behavior before L-DOPA exposure. Open-field testing showed no differences in total distance traveled (Fig. 3D) or velocity (Fig. 3E) between GFP- and sTREM2-injected mice. Lesion severity, assessed by apomorphine-induced rotations, was also unchanged (Fig. 3F), indicating that sTREM2 administered after lesioning neither improves nor worsens nigrostriatal injury. After LID induction, sTREM2 had no effect on locomotor activity, suggesting L-DOPA’s beneficial effects were not adversely affected by sTREM2 elevation (Fig. 3G). Compared with AAV-EGFP, AAV-sTREM2 resulted in a clear reduction in dyskinesia severity. sTREM2-injected mice showed significantly lower cumulative global dyskinesia scores (Fig. 3H), with reduced interval scores during weeks 2 and 3 (Fig. 3I). Thus, elevating sTREM2, an endogenous microglia-derived ligand, attenuates LID in a lesion model of Parkinson’s disease while leaving general locomotion and lesion severity unchanged.

To evaluate whether high sTREM2 levels interfere with the therapeutic benefit of L-DOPA, mice received sub-dyskinetic L-DOPA once weekly to avoid induction of dyskinesia (Fig. S7A,B). Open-field behavior and hemiparkinsonism-induced rotational bias were unchanged between groups at baseline and 0.75 mg/kg L-DOPA (Fig. S7C-F). At 1.0 mg/kg L-DOPA, sTREM2- and GFP-injected mice displayed similar increases in locomotion (Fig. S7G). Notably, at 1.0 mg/kg, sTREM2-injected mice also showed a significant reduction in rotational bias, consistent with preserved or potentially enhanced responsiveness to L-DOPA (Figure S7H). Cylinder testing further supported intact therapeutic L-DOPA effects. Baseline paw-use asymmetry was similar between groups, and sTREM2 did not alter rearing behavior. Sub-dyskinetic L-DOPA reduced paw-use bias in sTREM2-injected mice but not GFP controls (Fig. S7I-L), which may reflect testing prior to peak drug availability rather than true group differences. Nonetheless, across assays, sTREM2 did not diminish L-DOPA’s beneficial motor effects. Together, our results support sTREM2 as a microglia-derived modulator capable of improving dysregulated motor circuits without impairing dopaminergic benefit.

### Intravenous injection of AAV-sTREM2 partially reverses LID-associated SPN transcriptional changes

Microglia-derived signals regulate SPN pathway imbalance at the transcriptomic level during LID (Fig. 2). To test whether AAV-sTREM2 reverses these LID-associated transcriptional changes, we performed snRNA-seq on striatal nuclei from AAV-sTREM2- or GFP-treated mice harvested 1 h after the final L-DOPA injection during the on-state. Both lesioned and contralateral striata were analyzed (Fig. S8). A distinct LID-enriched dSPN subcluster (dSPN5; Fig. S9) emerged, closely matching the previously defined LID-responsive dSPN2 population (Fig. S6). Pseudobulk analysis comparing L-DOPA-treated sTREM2 and GFP groups showed downregulation of multiple immediate early genes (*Arc*, *Fosb*, *Junb*) in sTREM2-injected mice (Fig. 4A), paralleling the gene expression normalization observed after microglia repopulation. The sTREM2-induced changes aligned strongly with global LID-associated signatures: a striking 98% of sTREM2-regulated DEGs overlapped with LID-related genes (949/965 shared genes; Fig. 4B), and most were shifted in the opposite direction from LID induction (Fig. 4C). Because immediate early gene suppression (*Arc*, *Fosb*, *Junb*) suggested reduced dSPNs hyperactivity, we examined a curated activity-regulated gene set from active neurons in the dentate gyrus (*36*). A heatmap of averaged gene expression across dSPNs revealed an expected, stark increase in many activity-related genes in dSPNs exposed to L-DOPA (Fig. 4D). In comparison, co-exposure to sTREM2 blunted this response (Fig. 4D). Consistent with the transcriptomic results, c-Fos immunofluorescence showed that sTREM2 reduced the signal intensity of L-DOPA-evoked neuronal activation in the lesioned striatum (Fig. 4E-G).

**Fig. 4.**
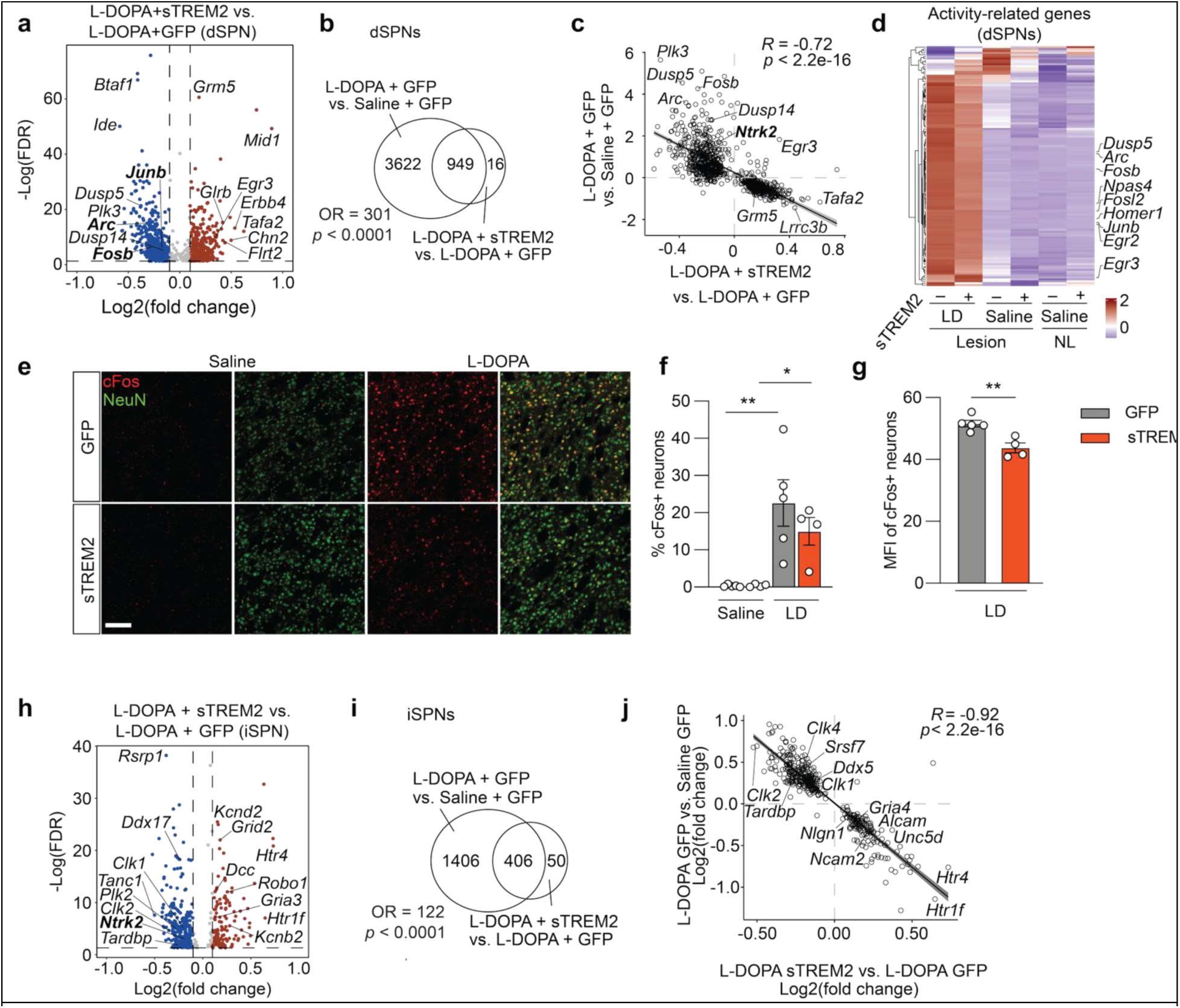
Intravenous injection of AAV-sTREM2 ameliorates LID-associated alterations in SPNs. **(a)** Volcano plot of dSPN DEGs comparing L-DOPA + sTREM2 versus L-DOPA + GFP. **(b)** Venn diagram of pseudobulk DEGs from GFP-injected mice treated with L-DOPA versus saline and from L-DOPA-treated mice injected with sTREM2 versus GFP. **(c)** Correlation of shared DEGs with regression and 95% confidence interval. Pearson correlation coefficients and p-values were determined with the ggpubr. **(d)** Heatmap of activity-related DEGs in dSPNs across conditions. **(e)** Representative cFos (red) and NeuN (green) immunofluorescence in lesioned dorsal striatum. Scale bar, 100μm. **(f)** Quantification of the percentage of cFos+ NeuN+ cells (mean ± s.e.m.). *p < 0.05, ***p < 0.001, repeated measures two-way ANOVA with Tukey’s test. **(g)** Mean fluorescence intensity of cFos+ NeuN+ cells in L-DOPA-treated GFP versus sTREM2 groups. **p < 0.01, unpaired t test. Each dot represents the average value across 4 sections from one mouse (n=5 saline-treated GFP-injected mice; n=4 saline-treated sTREM2-injected mice; n=5 L-DOPA-treated GFP-injected mice; n=4 L-DOPA-treated sTREM2-injected mice) **(h)** Volcano plot of iSPN DEGs comparing L-DOPA + sTREM2 versus L-DOPA + GFP. **(i)** Venn diagram of DEGs induced by L-DOPA (GFP) and DEGs altered by sTREM2 in L-DOPA-treated mice. **(j)** Correlation of shared iSPN DEGs between L-DOPA effects and sTREM2 effects. Pearson correlation coefficients and p-values were determined with the ggpubr. Dashed lines (a,h) show FDR < 0.05 and |log2 fold change| ≥ 0.1.

Elevation of sTREM2 also elicited distinct transcriptional changes in iSPNs. L-DOPA-treated iSPNs in sTREM2-injected mice showed decreased expression of RNA processing and splicing genes (*Ddx17, Clk1, Clk2, Tardbp*) and increased expression of ion channels and axon guidance molecules (*Grid2, Gria3, Kcnb2, Kcnd2, Dcc, Robo1*) (Fig. 4H). As in dSPNs, the sTREM2-induced changes in iSPNs aligned strongly with global LID-associated signatures: 89% of sTREM2-regulated DEGs overlapped with LID-related genes (406/456 shared genes; Fig. 4I). Many of these genes exhibited reversed expression with sTREM2 compared to the L-DOPA condition alone in iSPNs as well (Fig. 4J).

We further compared sTREM2-induced DEGs with those restored following microglia repopulation. dSPNs and iSPNs each showed significant overlap between sTREM2-responsive and microglia repopulation-responsive genes (Fig. S10A,B), with strong positive correlations in the direction and magnitude of change (Fig. S10C,D). Shared upregulated genes were enriched for synaptic pathways, whereas shared downregulated genes involved RNA splicing (Fig. S10E,F). dSPNs additionally showed enrichment of cytoskeletal pathways among downregulated terms, suggesting subtle divergence between SPN subtypes. Together, these findings show that AAV-sTREM2 counteracts a substantial fraction of LID-driven transcriptional alterations in both dSPNs and iSPNs. The convergence between sTREM2-induced and microglia repopulation-induced gene expression patterns supports a model in which microglia, in part through production of sTREM2, restrain dyskinesia-associated SPN hyperactivity and promote recovery of synaptic transcriptional programs.

### sTREM2 acts on TrkB and potentiates TrkB-ERK signaling

Given TrkB’s established role in synaptic plasticity (*37–39*), its altered expression following microglial depletion (Fig. 2A), and sTREM2 overexpression in SPNs of LID mice (Fig. 4H), we first asked whether sTREM2 directly interacts with TrkB. In vivo co-immunoprecipitation after AAV-sTREM2 delivery to wild-type mice revealed association of sTREM2 with both full-length and truncated TrkB (Fig. 5A). Consistently, split-luciferase complementation assays showed close proximity between sTREM2 and TrkB, which was diminished upon deletion of the TrkB extracellular domain (Fig. 5B,C), supporting a direct extracellular interaction.

**Fig. 5.**
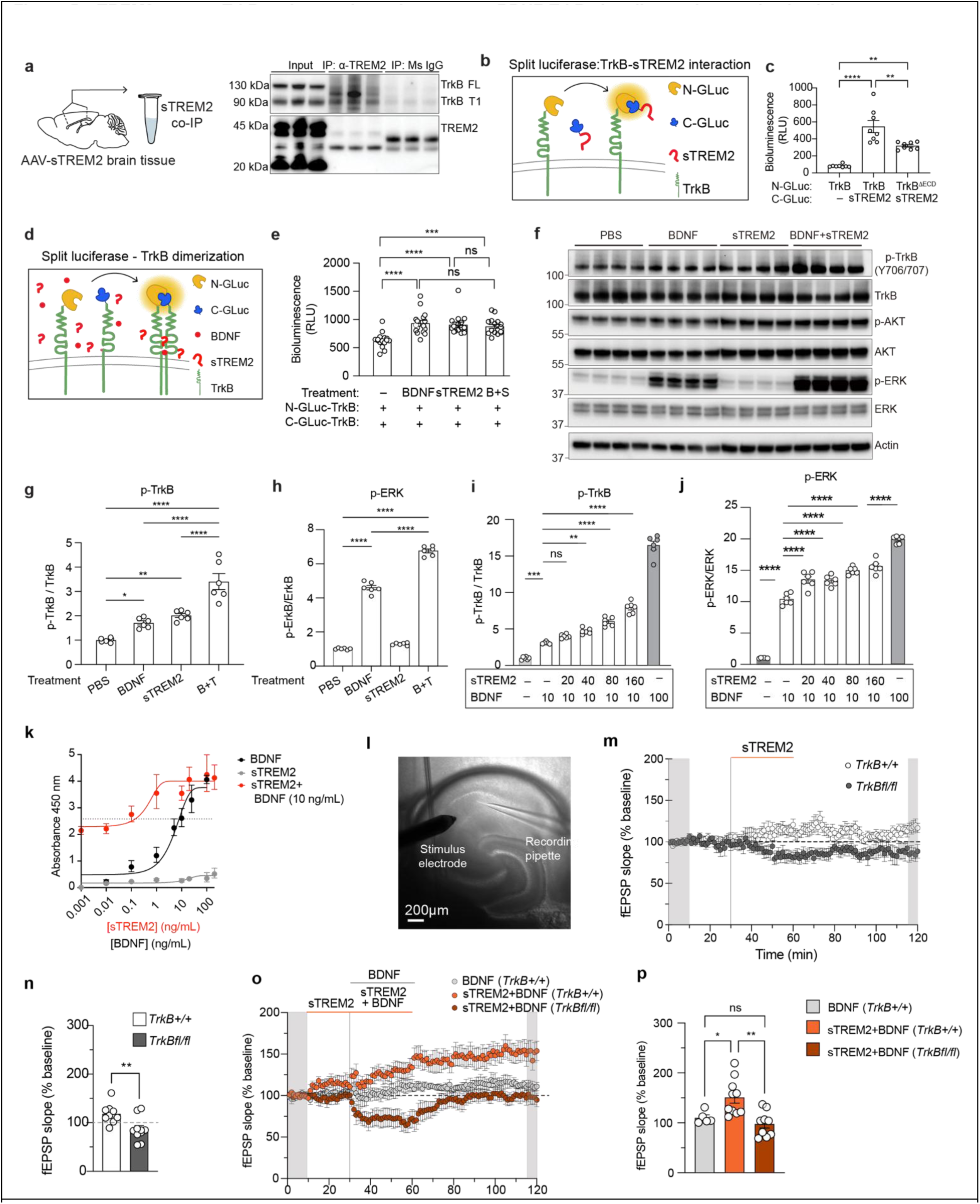
sTREM2 binds to TrkB and potentiates downstream BDNF-TrkB signaling and synaptic plasticity. **(a)** Schematic of cortical tissue collected from AAV-sTREM2-injected wild-type mice for co-IP (left). Immunoblot of TrkB and TREM2 after TREM2 or IgG pulldown (right; n = 3 mice). (**b)** Schematic of split luciferase assay testing sTREM2-TrkB interaction. (**c**) Quantification of bioluminescence from split luciferase assay for TrkB interaction (mean ± s.e.m.; n = 8 wells per condition). **p < 0.01**, \*\*\*\***p < 0.0001, one-way ANOVA with Tukey’s post hoc test. (**d**) Schematic of split luciferase assay testing TrkB dimerization. (**e**) Quantification of bioluminescence from split luciferase assay for TrkB dimerization (mean ± s.e.m.; n = 16 wells per condition). ***p < 0.001**, \*\*\*\***p < 0.0001, one-way ANOVA with Tukey’s post hoc test. (**f**) Representative lanes of Western blot for p-TrkB, TrkB, p-AKT, AKT, p-ERK, ERK, and actin in HEK 293T-TrkB^FL^ cells after treatment with BDNF (20ng/mL) and/or sTREM2 (200ng/mL) for 10 minutes. (**g–h**) Quantification of p-TrkB/TrkB (g) and pERK/ERK (h) (n = 6 samples per treatment). (**i–j**) Dose-dependent elevation of pTrkB/TrkB (**i**) and pERK/ERK (**j**) witih increasing concentrations of sTREM2 (10, 20, 40, 80, 160ng/ml) with 10 ng/ml BDNF, and 100 ng/ml BDNF alone (n = 6 samples per treatment). (**k**) ELISA measurement of p-TrkB induced by BDNF (0.001–100ng/ml), sTREM2 (0.001–100ng/ml) alone, or sTREM2 (0.001–100ng/ml) together with BDNF at 10 ng/ml. Each dot represents an average across 3-6 samples. (**l**) Representative hippocampal field recording configuration. (**m**) fEPSP slope time course with sTREM2 (200ng/mL) treatment. Each dot is the average (± s.e.m.) slope from all recorded slices (n = 9 or 10 slices from control or TrkB^fl/fl^ mice, respectively). Gray bars denote the time points averaged for baseline and post-treatment values. (**n**) Quantification of post-treatment fEPSP slopes relative to baseline. Each dot represents values from individual slices (n=1-2 slices per mouse; n = 5 or 8 mice/group). \*\**p* < 0.01, unpaired two-tailed t test. (**o**) fEPSP slope time course with BDNF (20ng/mL) and sTREM2 (200ng/mL) treatment. Each dot is the average (± s.e.m.) slope from all recorded slices (n = 5, 10, 9 slices from BDNF, sTREM2+BDNF, and sTREM2+BDNF in TrkB^fl/fl^, respectively). (**p**) Quantification of post-treatment fEPSP slopes relative to baseline. Each dot represents values from individual slices (n=1-2 slices per mouse; n = 5-7 mice/group). \**p* < 0.05, \*\**p* < 0.01, one-way ANOVA with Tukey’s post hoc multiple comparisons test.

We next examined whether sTREM2 modulates TrkB activation. Split-luciferase assays demonstrated that sTREM2 increased TrkB dimerization to a level comparable to low-dose BDNF (20ng/mL), with no additional enhancement upon co-treatment (Fig. 5D,E). In HEK 293T cells expressing full-length TrkB (HEK-TrkB-FL), sTREM2 increased TrkB phosphorylation, and this effect was further potentiated by BDNF (Fig. 5F,G). Downstream signaling analysis revealed pathway-biased modulation. BDNF robustly increased ERK phosphorylation, which was further elevated by sTREM2 (Fig. 5F,H). In contrast, although both BDNF and sTREM2 increased AKT phosphorylation without altering total AKT levels, no additive effect was observed at the level of p-AKT (Figure S11A,B).

To further define this interaction, HEK-TrkB-FL cells were treated with 10 ng/ml BDNF in the presence of increasing concentrations of sTREM2 (Fig. S11C). Quantification of p-TrkB/TrkB and p-ERK/ERK ratios showed that sTREM2 dose-dependently enhanced BDNF-induced TrkB and ERK phosphorylation (Fig. 5I,J). We next performed a highly sensitive p-TrkB ELISA using varying concentrations of sTREM2, BDNF, or their combinations (Fig. 5K). sTREM2 alone showed minimal intrinsic agonistic activity toward TrkB. However, in HEK-TrkB-FL cells treated with BDNF, co-application of increasing concentrations of sTREM2 produced a marked leftward shift in the BDNF dose–response curve, indicating enhanced ligand potency (reduced EC50). In addition, a modest increase in maximal p-TrkB levels was also observed at higher concentrations, suggesting that sTREM2 may slightly enhance signaling efficacy in addition to increasing BDNF sensitivity. Together, these results demonstrate that sTREM2 directly engages TrkB and potentiates BDNF-TrkB-ERK signaling. Meanwhile, sTREM2 acts as a positive allosteric modulator (PAM) of TrkB by enhancing both BDNF potency and maximal receptor activation.

### Acute sTREM2 enhances hippocampal synaptic plasticity via TrkB

To further establish the functional consequence of sTREM2-TrkB signaling, we turned to synaptic plasticity in the hippocampus, where TrkB signaling is a central regulator (*40–43*). Field excitatory postsynaptic potentials (fEPSPs) were recorded from the CA1 region in a manner consistent with exogenous BDNF application in previous studies (*44–48*) (Fig. 5L). Acute perfusion of hippocampal slices with sTREM2 (200 ng/ml) alone modestly increased fEPSP slopes relative to baseline (Fig. 5M,N). This increase was not observed upon conditional deletion of TrkB in excitatory CA1 neurons using local AAV-CaMKII-mCherry-Cre delivery in *TrkB^fl/fl^* mice (Fig. 5M,N), demonstrating that the effect of sTREM2 on hippocampal synaptic plasticity is TrkB dependent.

To determine whether sTREM2 modulates BDNF-TrkB signaling, hippocampal slices were perfused with a low concentration of BDNF (20 ng/mL) in the presence or absence of sTREM2 (200 ng/mL) (Fig. 5O). To minimize potential occlusion by BDNF during co-application with sTREM2, slices were pretreated with sTREM2 for 20 minutes. sTREM2 significantly potentiated BDNF-induced synaptic plasticity, and this enhancement was completely abolished by conditional deletion of TrkB in excitatory CA1 neurons (Fig. 5O,P). This effect is consistent with the ability of sTREM2 to potentiate BDNF-induced TrkB and ERK phosphorylation. Thus, sTREM2 enhances hippocampal synaptic plasticity by potentiating BDNF signaling in a TrkB-dependent manner.

### sTREM2 mimics the effects of BDNF on the dendritic excitability of SPNs

We next tested whether sTREM2 modulates striatal physiology in a manner consistent with its anti-dyskinetic effects through TrkB. Because the relative activity of dSPNs and iSPNs determines dyskinetic responses to levodopa (*30,49–51*)—with excessive dSPN activity and insufficient iSPN engagement contributing to dyskinesia (*52,53*)—we examined whether sTREM2 acutely alters cell-type–specific dendritic excitability under non-pathological conditions.

Backpropagating action potentials (bAPs) drive dendritic calcium influx and serve as instructive signals for synaptic plasticity (*54*). Using patch-clamp stimulation combined with calcium imaging, we quantified bAP-evoked calcium transients in proximal and distal dendrites of identified SPNs. Somatic stimulation reliably induced calcium signals in both compartments. Somatic stimulation reliably induced calcium signals in both compartments (Fig. 6A,B). In iSPNs, acute sTREM2 application significantly increased the amplitude of these calcium transients in both proximal and distal dendrites of iSPNs (Fig. 6C,D), indicating a broad increase in dendritic excitability. In contrast, sTREM2 only potentiated bAP-evoked calcium signals in proximal dendrites of dSPNs (Fig. 6E-G). Heat-inactivated sTREM2 had no effect, confirming dependence on biologically active sTREM2 (Fig. 6H,I).

**Fig. 6.**
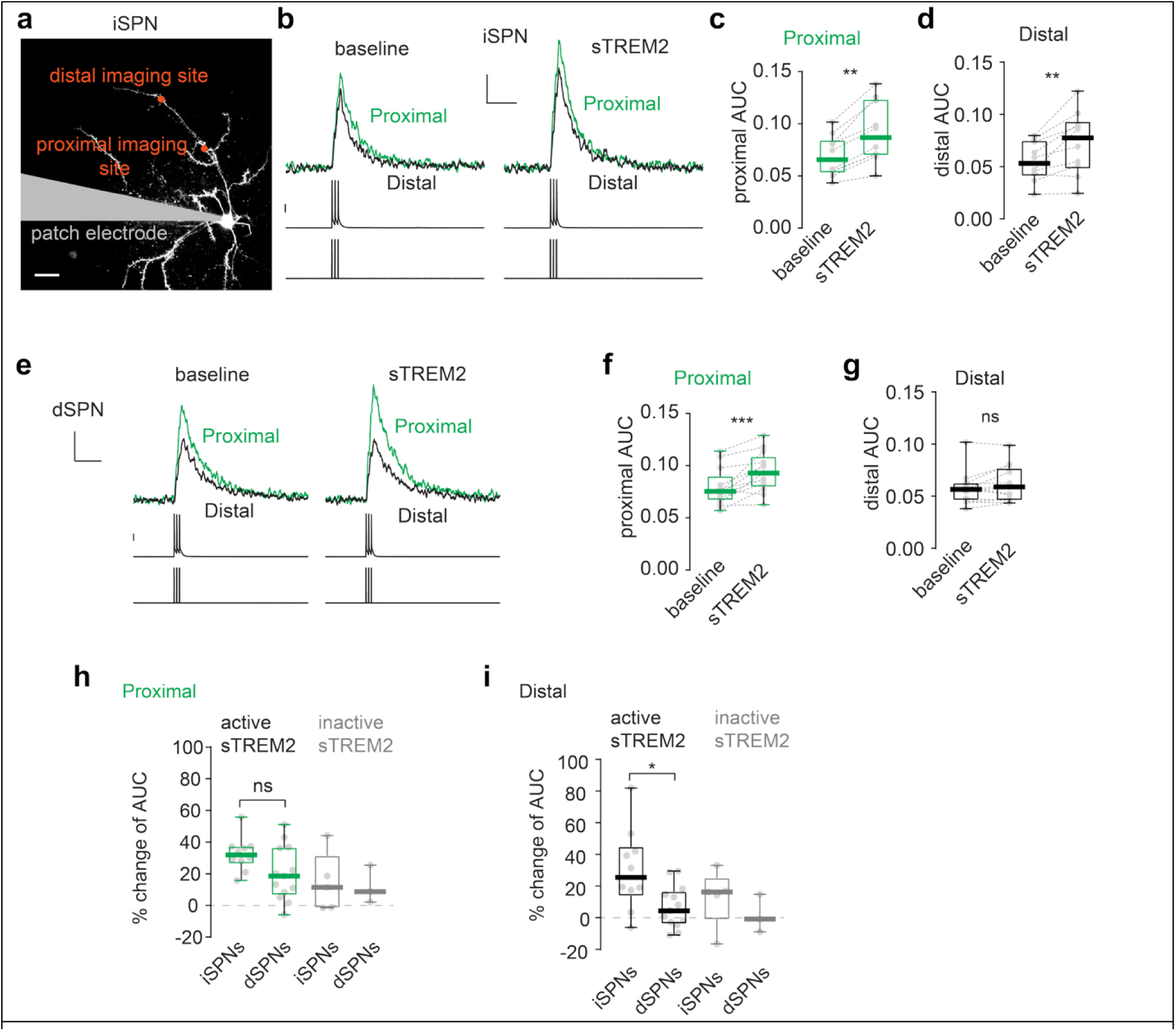
Acute sTREM2 application selectively increases dendritic excitability in SPNs. (**a**) Representative image of a whole-cell patched iSPN, with imaging sites on proximal and distal dendrites. Scale bar, 20μm. (**b**) Representative two-photon laser scanning microscopy (2PLSM) Fluo4 Ca^2+^ fluorescence measurements at proximal and distal dendrites of an iSPN evoked by brief somatic current pulses (2nA, 2ms, 20 msec inter-stimulus interval; bottom); somatic voltage response showing spike generation is shown at the bottom (scale bar, 25mV; middle). Dendritic Ca^2+^ signals evoked before and after sTREM2 (concentration) application (scale bars, 0.1 ΔG/R_0_ and 0.2s). (**c–d**) Summary of proximal (**c**) and distal (**d**) dendritic Ca^2+^ signals (AUC of ΔG/R_0_) of SPNs before and after acute sTREM2 application (n=10 cells from 10 mice). (**e**) Representative recordings from a dSPN showing the fluorescence transients evoked in proximal and distal dendrites, respectively. Scale bars are as described for **b**. (**f–g**) Summary of proximal (**f**) and distal (**g**) dendritic excitability (AUC of ΔG/R_0_) in dSPNs before and after acute sTREM2 application (n=13 cells from 10 mice). Data are presented as non-parametric box-and-whisker plots. (**h–i**) Summary of changes in dendritic excitability at proximal (**h**) and distal (**i**) dendrites of iSPNs and dSPNs following acute application of sTREM2 or heat-inactivated sTREM2 (for heat-inactivated sTREM2, n=5 iSPNs from 4 mice and n=3 dSPNs from 3 mice). Data are presented as non-parametric box-and-whisker plots. For c, d, f, and g, **p < 0.01, ***p < 0.001, paired Wilcoxon signed-rank test. For h and i, *p < 0.05, unpaired Mann-Whitney test.

These findings align with prior work showing that chemogenetic restoration of balanced excitability in iSPNs and dSPNs—similar to the effect produced by sTREM2—attenuates LID (*52*). By preferentially increasing dendritic responsiveness in iSPNs while avoiding excessive distal amplification in dSPNs, sTREM2 recapitulates key features of BDNF–TrkB signaling in a cell type–specific manner and rebalances activity between the indirect and direct pathways.

## Discussion

In this study, we identify a neuroimmune mechanism by which microglia restrain maladaptive striatal plasticity underlying L-DOPA–induced dyskinesia (LID). We show that microglia exert a protective influence during dyskinesia and that elevating the microglia-derived ligand sTREM2 alleviates abnormal involuntary movements without compromising the therapeutic benefit of L-DOPA. Acute sTREM2 enhances dendritic excitability of striatal spiny projection neurons in a compartment- and cell type-specific manner, consistent with a circuit-level rebalancing of striatal output, recapitulating key features of BDNF–TrkB signaling in counteracting LID. Mechanistically, sTREM2 directly engages TrkB, promotes TrkB phosphorylation, and dose-dependently potentiates BDNF-driven downstream signaling in a TrkB-dependent manner. These findings redefined sTREM2 as a positive allosteric modulator of BDNF-TrkB, tuning activity-dependent plasticity beyond a classical inflammatory cue alone.

While microglial depletion using CSF1R inhibitors is a coarse intervention whose impact is highly dependent on disease stage, our microglial depletion and repopulation experiments further clarify the role of microglia in LID—a common feature of late-stage PD. In toxin-based models, prior studies have shown that microglial depletion before lesioning dopaminergic neurons exacerbates their loss and subsequent motor deficits (*55–57*). Here, we depleted microglia after establishing the parkinsonian state, at the time of LID induction. This late intervention markedly worsened dyskinesia, which was reversed by microglial repopulation. Thus, microglia exert an adaptive and protective effect across multiple stages of PD, and their loss aggravates disease-related striatal dysfunction.

Maladaptive striatal plasticity is a core feature of LID. Imbalanced activity of ‘pro-movement’ dSPNs and ‘anti-movement’ iSPNs after sustained striatal dopamine concentration plays a critical roles in LID. Chemogenetic rebalancing of dSPN and iSPN excitability alleviates LID (*52*). By acting through TrkB, sTREM2 appears to achieve similar results. Previous work established that BDNF signaling through TrkB increases the dendritic excitability of iSPNs (*51*), as did sTREM2. This was accompanied by a more modest effect in dSPNs that was limited to proximal dendrites. Both BDNF and TrkB are elevated in the striatum of LID models and striatal deletion of TrkB worsened LID (*30,58*). Although our dendritic excitability results highlight iSPNs in mediating the anti-dyskinetic effects of sTREM2, chronic sTREM2 effects likely involve dSPNs, as evidenced by the robust transcriptomic changes in both SPN subtypes (*49,59,60*).

A central advance of this study is the discovery that sTREM2 acts as a putative endogenous positive allosteric modulator of BDNF. Co-immunoprecipitation and split-luciferase complementation assays demonstrate that sTREM2 directly engages the TrkB extracellular domain, promotes receptor dimerization, and enhances TrkB phosphorylation. sTREM2 further dose dependently potentiates TrkB and ERK activation induced by 10 ng/ml BDNF. Ultrasensitive biosensor measurements indicate that perisynaptic BDNF can transiently rise into the low ng/ml range in vivo (*61*). This dynamic range overlaps with the concentration at which sTREM2 enhances TrkB signaling, supporting a physiologically relevant modulatory role. Notably, sTREM2 did not augment AKT phosphorylation in TrkB expressing HEK cells, suggesting selective bias toward ERK dependent plasticity pathways. Important mechanistic questions remain. It is unclear which domains of sTREM2 bind to TrkB or whether the interaction stabilizes ligand bound receptor complexes, alters receptor conformation, or modulates TrkB membrane organization and trafficking. These mechanisms will require further biochemical and structural investigation.

In further support of sTREM2’s potentiatiation of TrkB signaling, we find that sTREM2 enhances hippocampal BDNF-induced synaptic plasticity in a TrkB-dependent manner. Acute sTREM2 robustly amplifies BDNF-induced long-term potentiation, effects that are abolished by conditional deletion of TrkB in excitatory CA1 neurons. These findings extend prior observations that high concentrations of sTREM2 enhance hippocampal plasticity (*62,63*) and establish TrkB as a critical mediator of this effect. Together with our striatal data, these results indicate that sTREM2 broadly tunes TrkB-dependent plasticity across circuits implicated in dyskinesia and cognition, suggesting a generalized role for sTREM2 in regulating activity-dependent neuronal adaptation.

We provide evidence for AAV-sTREM2 as a potential therapeutic for LID that does not interfere with the therapeutic efficacy of L-DOPA. Studies on sTREM2 have mainly focused on its use as a biomarker in cerebrospinal fluid (CSF) for neurodegenerative disease. The ability of sTREM2 to amplify endogenous BDNF within its physiological dynamic range enables microglia to fine-tune synaptic plasticity with the release of sTREM2, which is regulated by genetic risk factors and microglial states. With the exception of a rare allele that causes near complete loss of membrane bound TREM2 (*19,62*), genetic variants associated with higher cerebrospinal fluid sTREM2 levels are most often linked to lower disease risk (*64*), supporting protective roles for both membrane bound and soluble TREM2. AAV-mediated sTREM2 overexpression represents a unique way to enhance the protective effects of microglia while preserving endogenous microglia TREM2 signaling. Consistent with this, experimental elevation of sTREM2 ameliorates pathology in amyloid and tau models (*63,65,66*). AAV-sTREM2 has been reported to mitigate several aspects of AD pathology, like amyloid plaque load (*54*) and tau hyperphosphorylation via binding to transgelin 2 (*63,65*). Here, we broaden the therapeutic potential of AAV-sTREM2 by showing an amelioration of LID both behaviorally and transcriptomically. Whether earlier administration of AAV-sTREM2 can slow or attenuate dopaminergic cell death in this model or other PD models that account for alpha-synuclein pathology remains a subject for future work.

In summary, we propose that microglia protect against LID by releasing sTREM2, which engages and potentiates neuronal TrkB signaling to restrain maladaptive plasticity. By enhancing TrkB-dependent pathways, sTREM2 promotes balanced dendritic excitability and, in principal, synaptic remodeling in striatal and hippocampal circuits. More broadly, our findings identify sTREM2 as a concentration-dependent, microglia-derived neuromodulator and a potential endogenous positive allosteric modulator of TrkB. Rather than serving as a binary inflammatory cue, sTREM2 scales TrkB-ERK signaling across a graded range, enabling microglia to tune synaptic strength and circuit function. This work redefines a neuroimmune mechanism through which microglial state dynamically shapes neuronal adaptability and synaptic plasticity.

## Acknowledgments

We would like to thank Dr. Tara Tracy (Buck Institute) for helpful discussions; Drs. Chunyu Wang (Rensselaer Polytechnic Institute) and Kevin Nash (University of South Florida) for advice on split-luciferase assays; Guillermo Coronas and Ana Carney for administrative support, and Weill Cornell Medicine’s Microscopy and Imaging Analysis Core.

## Funding

This work was supported by:

Freedom Together Foundation (to LG, MGK, and DJS)

National Institutes of Health grant R01AG076448 (to LG)

National Institutes of Health grant R01AG072758, R01AG079557-01, and 1R01AG079291-01A1 (to LG)

National Institutes of Health grant R01AG074541 (to LG)

National Institutes of Health grant R01NS126590 (to FSL)

National Institutes of Health grant RF1AG07861 (to FSL)

Rainwater Charitable Foundation (to LG)

Cure Alzheimer’s Fund (to LG)

Pritzker Neuropsychiatric Disorders Research Consortium (to FSL)

## Author contributions

Conceptualization: LG, MGK, CC, RM, YW, ERT

Data curation: LF, CC, LG, QC

Funding acquisition: LG, MGK, DJS, FSL

Investigation: CC, RM, YW, ERST, LF, MWJ, QC, JK, PY, JZ, AP, ABF, KN, BG, NF, MYW, DZ, MB

Methodology: CC, RM, YW, ERST, MWJ, QC, JK, PY, MYW, SG

Software: CC, LF, MWJ, QC, ERST

Supervision: LG, MGK, DJS, FSL, AGO

Visualization: CC, YW, ERST, LF, MWJ, QC, JK, NF, MYW

Writing—original draft: CC, LG

Writing—reviewing and editing: LG, CC, DJS, FSL, RM, YW, ABF

## Data, code, and materials availability

Previously published bulk RNA sequencing data from human PD patients (*23*) can be accessed through the GEO database (GEO205450). Previously published microarray data from the striatum of LID mice (*27*) can be accessed through the GEO database (GSE55096). Single nucleus RNA sequencing from activated neurons in the hippocampus of mice (*36*) can be accessed through the GEO database (GSE77067). Single nucleus RNA sequencing data generated in this study were deposited into the GEO database under accession number GSE324025 will be made publicly available upon publication. Original code for snRNA-seq analysis can be found at: https://github.com/lifan36/Castagnola-Marongiu-Wan-LID-2026. Code for calcium imaging analysis can be found at: https://doi.org/10.5281/zenodo.17380534. All materials used in this study can be obtained upon request to L.G.

## Supplementary Materials

### Materials and Methods

#### Weighted correlation network analysis (WGCNA) on basal ganglia tissue from PD and control patients

Bulk RNA-sequencing data and corresponding metadata from the brain of PD patients or healthy controls from a previously published paper (*23*) were accessed through the GEO database (GEO205450). Genes with too many missing values were detected using the *goodSamplesGenes* function in the WGCNA (*67*) R package (v1.73) and removed from further analysis (n = 189 genes). The size factors and dispersion estimates for the count matrix was performed using the default *DESeq* function in the DESeq2 package (v1.42.1). The count matrix was then transformed using vst in the DESeq2 package. To constrain the amount of genes for downstream analysis, only the top 5% of the genes in the transformed data were used for WGCNA analysis. The WGCNA network was constructed using the *blockwiseModules* function with the power set to 9 (generated via the *pickSoftThreshold* function), a ‘signed’ network, and a minimum module size of 30. Module labels were assigned to colors and dendrograms for clustered gene assignment were generated using the WGCNA package. Genes in each module were extracted and the module eigengene values was retrieved with the WGCNA package. The gray module, which contains genes not assigned to any other module, was discarded. Dendrograms and heatmaps of eigengenes were constructed with the *plotEigengeneNetworks* function in WGCNA. To correlate metadata traits with module eigengenes, categorical data was binarized and control or unknown values were removed from PD-related traits (i.e. dementia, dyskinesia, onset age, and cholinesterase inhibitors). Pearson correlation coefficients between eigengenes and traits were obtained using the *cor* function in the stats package (v4.3.2). P-values for the module-trait correlations were determined with the *corPvalueStudent* function in the WGCNA package then adjusted using the Benjamin-Hochberg procedure (FDR). Heatmaps of module-trait correlations were generated with the *labeledHeatmap* function. The signed adjacency (or correlation) matrix for genes in the black module was obtained with the *adjacency* function using the same power as the network generation, 9, and the corresponding heatmap was generated with the ComplexHeatmap (v2.18.0) and circlize (v0.4.16) packages. To generate the network graph of genes in the black module, a topological overlap matrix (TOM) was generated from the gene expression for the corresponding genes with the *TOMsimilarity* function. This node-edge list was uploaded into the Cytoscape application (v3.10.3) for creating the network graph with edge values below 0.15 filtered out. Module preservation analysis is outlined in the single nuclei RNA sequencing section below.

#### Pathway analysis of gene sets

Gene symbols were converted to their Entrez ID values using the AnnotationDbi package (v1.64.1) in R with the appropriate species gene annotation database, org.Hs.eg.db (v3.18.0) or org.Mm.eg.db (v3.18.0). Over-representation analysis in the KEGG or GO databases was performed with the *enrichKEGG* or *enrichGO* functions in the clusterProfiler package (v4.10.1). For GO analysis, all ontology categories were probed and redundant terms (with a similarity cutoff of 0.5) were removed from the output. Bubble plots were generated with ggplot2 (v3.5.2).

#### Mice

All experiments were performed in accordance with NIH guidelines and approved by the Institutional Animal Care and Use Committee at Weill Cornell Medicine and Northwestern University. Mice were group housed in temperature- and humidity-controlled rooms on a 12-h light/12-h dark cycle with food and water available *ad libitum.* For 6-OHDA experiments, two-month-old male wild-type C57BL/6J mice were purchased from Jax (strain #000664). All behavior tests were performed during the light-on cycle. For *ex vivo* calcium imaging experiments, male C57BL/6 mice (9-14 weeks old) hemizygous for tdTomato or eGFP expression under control of *Drd1a* or *Drd2* regulatory elements (RRID: MMRRC_030512-UNC and RRID: MMRRC_000230-UNC, backcrossed to a C57BL/6 background) were used. *TrkB^fl/fl^* mice were kindly provided by Dr. Baoji Xu at University of Florida. For *ex vivo* hippocampal recordings, male and female wild-type C57BL/6J, *TrkB^fl/fl^* and their *TrkB^+/+^* littermate controls (2-4 months old) generated from heterzygous matings were used.

#### Unilateral 6-hydroxydopamine (6-OHDA) lesioning

5-6 month-old male mice were anesthetized with ketamine (110 mg/kg)/xylazine (4.4-10 mg/kg; i.p.), given carprofen (5mg/kg; s.c.), and pre-treated with desipramine (25 mg/kg; i.p.). Mice were head-fixed in a stereotactic frame (Kopf Instruments) and injected unilaterally with 600-800 nL of 6-OHDA (3mg/mL solution) delivered at 100 nL/min into the left medial forebrain bundle (AP: −1.1mm, ML: −1.1mm, DV: −5.0mm, relative to bregma). 6-OHDA injections were started 30 minutes after administration of desipramine to prevent toxicity to noradrenergic neurons. Carprofen (5 mg/kg; s.c.) was delivered again 24 hours after surgery. After surgery, mice received twice daily injections (s.c.) of 1mL Lactate Ringer’s solution containing 5% Dextrose solution and were given condensed milk, peanut butter cups, and moistened food pellets every day at the bottom of their cage for 2 weeks (or until fully recovered) to prevent dehydration and excessive weight loss. Mice were monitored for signs of distress, inflammation, and/or infections.

#### Hippocampal CA1 viral injections

*TrkB^fl/fl^* or *TrkB^+/+^*mice were anesthetized with isoflurane and given meloxicam (2mg/kg; s.c.). Mice were head-fixed in a stereotaxic frame (RWD) and injected bilaterally with 250nL of 3.5E12 GC/mL AAV5-CaMKII-mCherry-Cre (UNC Vector Core; Lot No. AV6448D) into the CA1 (−2.2mm AP, +/− 1.5mm ML, and −1.25mm DV from bregma) at 100nL/min. Recordings were performed 3-4 weeks following injections.

#### Apormorphine-induced rotation test for behavioral confirmation of striatal lesions

Five weeks after 6-OHDA lesions, mice were given 0.25 mg/kg apomorphine (s.c.) and placed into circular open field chambers. Locomotion was recorded and tracked for 30 minutes using EthoVision XT and the net number of contralateral turns to the injection site (i.e. clockwise turns) was used to determine lesion efficacy. Mice that exhibited over 3 turns/min were considered successfully lesioned. Mice that did not reach this threshold were re-lesioned with 6-OHDA, reassessed, and excluded from further study if they failed to reach the threshold after the second rotation test. Successfully re-lesioned mice showed similar net contralateral turns compared to mice that only received one 6-OHDA lesion surgery. Mice that recovered from the initial lesion, that is, met the threshold during the first rotation test but did not meet threshold during the second rotation test, were also excluded from further study. After all apomorphine-induced rotation tests, further behavioral assays were performed at least one month after to reduce priming effects from apomorphine, a dopamine receptor agonist.

For sTREM2 overexpression studies, rotation tests were performed again six weeks after lateral tail vein injections to assess if sTREM2 overexpression alters the locomotor response to apomorphine.

#### Lateral tail vein injections for viral delivery

Mice are warmed under a heat lamp to induce venodilation then placed into a restraint device that provides access to the tail. The tail is cleaned with a sterile alcohol wipe. 100μL of either AAV-PHP.eB-CAG-sTREM2-T2A-GFP or AAV-PHP.eB-CAG-GFP (1E11 GC/mL) was slowly injected into the lateral tail vein. Pressure and gauze were applied to the injection site to stem any bleeding then mice were returned to their home cage.

#### Open field test for general locomotion

Six weeks after lateral tail vein injections, mice were habituated to the behavior room for 1 hour before handling and to the open field chambers for 10 minutes each day for two days prior to testing. On the testing day, mice were brought to the behavior room one hour before testing started. Locomotion was recorded and tracked for 30 minutes using EthoVision XT and the total distance traveled, average velocity, and net ipsilateral turns (i.e. counter-clockwise turns) was assessed.

#### Cylinder test to assess forepaw use bias

Mice were habituated to the room for one hour before beginning the test. Mice were injected with L-DOPA or saline five minutes before testing. During the test, individual mice were placed in a glass cylinder in front of two angled mirrors to track paw usage at all times. Mice were video recorded for 3 minutes. The mouse undergoing testing was not able to observe other mice or the tester during the test. After the test, mice were returned to the home cage and the cylinder was cleaned with 70% ethanol in between tests or between each cage of mice. The number of paw contacts made with the side of the glass cylinder for each paw and the time spent rearing, excluding grooming, was scored offline using BORIS (v9.3.2). The examiner was blinded during the scoring.

#### Abnormal involuntary movements (AIMs) scoring

AIMs were scored as previously published (*68*). Briefly, on the day of testing, mice were habituated to clear Plexiglas containers spaced at least 10cm apart on a freestanding metal rack for at least one hour. Each mouse was scored for 1 minute every 20 minutes following L-DOPA injections (s.c.) until dyskinesias were no longer observable. Subtypes of dyskinesia (axial, limb, and orolingual) were monitored by a blinded examiner continuously across that minute and were scored according to the frequency (basic) and severity (amplitude) of each dyskinesia subtype. Basic and amplitude scores for each subtype of dyskinesia were multiplied to obtain global dyskinesia subtype scores. Global scores for each subtype were added together for the global dyskinesia score and the cumulative global dyskinesia score represents the summed global dyskinesia scores across the scoring session. A similar scale for non-dyskinetic locomotion was used during scoring.

#### Perfusions and tissue collection

Mice were injected with the final dose of saline or L-DOPA one hour before euthanization with FatalPlus. Blood was collected via cardiac puncture then the mouse was perfused intracardially with PBS kept on ice. Blood was spun down at 5000rpm for 5 minutes and plasma was frozen at −80°C until used. After perfusion, a small portion of the posterior cortex and midbrain was dissected from all mice for post-mortem sTREM2 overexpression confirmation. Hemispheres reserved for immunofluorescence were post-fixed in 4% PFA overnight at 4°C with gentle agitation. The remaining tissue was microdissected with a 0.5mm coronal mouse brain matrix and fine forceps to extract the cortex, hippocampus, striatum, substantia nigra, and thalamus. Microdissected regions were frozen on dry ice immediately after dissection and stored at −80°C until further use.

#### Single nuclei RNA sequencing

Nuclei isolation from frozen striatal tissue was adapted from a previous study (*69*), with modifications. Frozen striatal tissue was placed in 1500μL of nuclei PURE lysis buffer (Sigma-Aldrich; Cat. No. NUC201) and homogenized with a Dounce tissue grinder (Sigma-Aldrich; Cat. No. D8938) using 15 strokes per pestle. Homogenized tissue was filtered through a 35μm cell strainer, centrifuged at 600g for 5 minutes at 4°C then washed three times with 1mL of PBS containing 1% BSA (Thermo Fisher Scientific; Cat. No. 37525), 20mM DTT (Thermo Fisher Scientific; Cat. No. 426380500), and 0.2 U/μL recombinant Ambion RNase inhibitor (Invitrogen; Cat. No. AM2684). Nuclei were centrifuged at 600g for 5 minutes at 4°C then resuspended in 350μL of PBS containing 0.04% BSA and 1xDAPI followed by fluorescence-activated cell sorting to remove cell debris. The DAPI-stained nuclei in the sorted suspension were counted and diluted to 1000 nuclei/μL in PBS containing 0.04% BSA. All procedures were performed on ice or at 4°C.

For droplet-based snRNA-sequencing, libraries were prepared with Chromium Single Cell 3’ Reagents Kits (v3.1; 10x Genomics; Cat. No. PN-1000075) according to the manufacturer’s instructions. The snRNA-seq libraries were sequenced on a NovaSeq 6000 sequencer (Illumina) with PE 2x50 paired-end kits by using the following read length: 28 cycles Read 1, 10 cycles i7 index, 10 cycles i5 index, and 90 cycles Read 2.

Reads were aligned to the mm10 genome using Cell Ranger (10x Genomics; v6.1.2) to obtain gene counts. Reads mapped to pre-mRNA were counted. Default parameters in Cell Ranger were used to call cell barcodes. Genes detected in three or fewer cells were excluded from downstream analyses. Cells were filtered based on gene complexity, total UMI counts, and mitochondrial read content, with quality-control thresholds tailored to each dataset. For the LID dataset, cells were retained if they had 300–4,000 detected genes (nFeature_RNA), fewer than 20,000 UMIs (nCount_RNA), and <5% mitochondrial reads (percent.mt). For the LID_PLX dataset, cells were retained if they had 300–5,000 detected genes, fewer than 15,000 UMIs, and <1% mitochondrial reads. For the LID_sTREM2 dataset, cells were retained if they had 300–6,000 detected genes, fewer than 35,000 UMIs, and <5% mitochondrial reads. The DoubletFinder R package was used to predict potential doublet cells for each sample separately and high confidence doublets were removed (*70*). Count normalization and clustering was performed with the Seurat package (v3.2.2). Briefly, counts for all nuclei were scaled by the library size, multiplied by a scale factor of 10000, and log-transformed. The top 2000 highly variable genes were identified using the *FindVariableFeatures* function in Seurat with the vst method on SCTransform data from the sctransform package. This returned a corrected UMI count matrix, a log-transformed data matrix, and Pearson residuals from the regularized negative binomial regression model. PCA analysis was done on all genes, and t-SNE was run on the top 15 PCs. Cell clusters were identified with the Seurat functions *FindNeighbors* (dims = 1:15) and *FindClusters* (resolution = 0.1). The neighborhood size parameter (pK) was estimated using BCmvn with 15 PCs and the pN set to 0.25 by default. Sample integration was performed using the Seurat functions *FindIntegrationAnchors* and *IntegrateData*.

For snRNA-seq of lesioned and non-lesioned striatal tissue (Figure 1 and Extended Figure 3), we used saline- (n=5) or L-DOPA-treated (n=5) mice and collected the striatum from both the lesioned and non-lesioned hemispheres in each mouse (n=20 samples total). For snRNA-seq of lesioned striatal tissue from the microglia depletion experiment (Figure 2.5 and 2.6), the striatum from the lesioned hemisphere was collected from saline-treated control diet mice (n=3), L-DOPA-treated control diet mice (n=3),L-DOPA-treated PLX diet mice (n=3), L-DOPA-treated PLX reversal diet mice (n=3), and acute L-DOPA-treated mice (n=3). Acute L-DOPA-treated mice were administered saline for the duration of the experiment but received one injection of L-DOPA prior to perfusion. For snRNA-seq of lesioned and non-lesioned hemispheres from the sTREM2 overexpression experiment (Figure 4 and Extended Figure 8), the striatum from the lesioned hemisphere of L-DOPA-treated GFP-injected (n=4) and L-DOPA-treated sTREM2-injected (n=4) mice and the striatum from the lesioned and non-lesioned hemisphere of saline-treated GFP-injected (n=4) and saline-treated sTREM2-injected (n=4) mice were collected.

Each cluster was assigned a cell type label by using statistical enrichment for sets of marker genes in each cluster with manual evaluation for known marker genes. Subclustering of each cell type was performed by subsetting the data set, re-normalizing the counts, and re-clustering each cell type with the method outlined above, parameters for the *FindNeighbors* and *FindClusters* functions were tailored for each cell type as needed. DEG analysis was performed for each cell type with the *FindMarkers* function in Seurat using MAST (*71*).

To assess human gene module preservation in mouse striatal microlia (Figure 1 and Extended Figure 4), we first converted human genes to the corresponding mouse orthologs with the orthogene package (v1.8.0). Genes not mapped to a mouse ortholog or not detected in any microglia were dropped from further module preservation analysis. At least 50% of the orthologous genes in each module were detected in mouse microglia. Preservation analysis was performed using the hdWGCNA (*26*) package (v0.4.06) which leverages the *modulePreservation* function in the WGCNA package for calculation of a Z statistic. As described in the original module preservation paper (*25*), the Z statistic for each module is determined by comparing the observed module assignment to random module assignment. Z values greater than 2 are considered moderately preserved and values greater than 10 are considered strongly preserved.

### Brain tissue protein assay

Microdissected cortical tissue was thawed on ice and weighed. RIPA buffer (Thermo Fisher Scientific; Cat. No. 89901) with 1X cOmplete Protease Inhibitors (Roche; Cat. No.11697498001) and 1X Halt Phosphatase Inhibitors (Thermo Fisher Scientific; Cat. No. 1862495) was added to the tissue (1:10 w:v). Samples were sonicated (Epigentek; Model EQC-2000) for 5 minutes at 40% amplitude with 5 seconds-on and 5 seconds-off pulses until there were no visible tissue chunks remaining. Samples were then centrifuged at 20000g for 15 minutes at 4°C and the supernatant was collected. Supernatant diluted in 1:15 ddH_2_O in a separate tube. Supernatant was stored at −80°C until further use. The protein concentration was determined with the Pierce BCA protein assay (Thermo Fisher Scientific; Cat. No. 23225). Albumin standard was made using serial dilutions from 0-2mg/mL. Diluted samples or albumin standard (10μL) was added to each well of a 96-well plate, run in duplicate. BCA reagent A was mixed with BCA reagent B (50:1 respectively) and 200μL was added to each well immediately after the mixture was prepared. Samples were incubated at 37°C for 30 minutes then absorbance was read at 562nm on a plate reader (BioTek; Synergy H1 Hybrid Reader). Protein concentration was estimated using a standard curve and averaged absorbance values for each sample.

### Automated sandwich ELISA for TREM2

Sample supernatant from cortical lysate (20μg) was diluted 1:5 in sample diluent and 50μL was added to each well of the TREM-2 cartridge (biotechne; Cat. No. SPCKB-PS-001847) with 1mL of wash buffer. For plasma values, 14μL of plasma was diluted 1:5 in sample diluent and 50μL was added to each well. The film on the bottom of the cartridge was removed and the cartridge was placed into an automated immunoassay system (v3.7.2.0; Protein Simple; Cat. No. 600-100). TREM2 concentration was estimated using a standard curve with averaged replicate values for each sample. Data was input into GraphPad Prism to generate an ROC curve, the threshold for inclusion in the sTREM2 overexpression group was based on the value closest to the top-left corner of the curve. Mice with brain tissue TREM2 concentration values that did not meet this threshold were excluded.

### Immunofluorescence, imaging, and analysis

Following overnight post-fixation at 4°C, hemibrains were transferred to 1XPBS with 0.1% sodium azide. Before sectioning, brains were placed in 30% sucrose in deionized water with 0.1% sodium azide at 4°C for at least 3 days. Brains were sectioned coronally into 30μm sections on a sliding microtome (Leica; Model SM2010R) set to −28°C initially then raised to −23°C once the brain was frozen. Sections were collected to 1.5mL Eppendorf tubes containing cryoprotectant solution (F.D. Neurotechnologies; Cat. No. PC101) and stored at −20°C.

Striatal sections (n=4 sections per mouse, n=4-5 mice per condition) were transferred to a 24-well plate (2 sections per well) and washed in 1XPBS (gibco; Cat. No. 70011-044) 3 times for 10 minutes, shaking at room temperature. Sections were incubated in 0.5% Triton X-100 () in PBS (PBST) for 10 minutes, shaking at room temperature, then washed twice in 1XPBS for 5 minutes. Sections were then blocked for 30 minutes in 10% normal goat serum (Jackson Labs; Cat. No. 005-000-121) in PBS and washed twice in 1XPBS for 5 minutes on a shaker. Sections were blocked in mouse-on-mouse mouse Ig blocking reagent (Vector Laboratories; Cat. No. BMK-2202) for 1 hour shaking at room temperature then washed twice in 1XPBS for 5 minutes. Sections were then incubated with primary antibodies diluted in M.O.M diluent (Vector Laboratories; Cat. No. BMK-2202) overnight shaking at 4°C. The following day, sections were washed in 1XPBS three times, shaking for 5 minutes at room temperature. Secondary antibodies were diluted in M.O.M. diluent and added to the sections and incubated for 1 hour shaking at room temperature, shielded from light. Sections were washed in 1XPBS three times for 5 minutes then mounted onto slides. Slides were left to dry then coverslipped with ProLong Gold Antifade mounting media (Invitrogen; Cat. No. P36930). Primary antibody dilutions and information are as follows: guinea pig anti-NeuN (Millipore; Cat. No. ABN90) at 1:500; rabbit anti-cFos clone 9F6 (Cell Signaling; Cat. No. 2250S) at 1:500; rabbit anti-Iba1 (Wako; Cat. No. 019-19741) at 1:250; chicken anti-tyrosine hydroxylase (Abcam; Cat. No. ab76442) at 1:500. Sections used for negative controls were only incubated with NeuN primary antibody. Secondary antibody dilutions and information are as follows: 488 goat anti-Guinea pig (Invitrogen; Cat. No. A11073) at 1:500; 647 goat anti-rabbit (Invitrogen; Cat. No. A32733) at 1:500; 568 goat anti-rabbit (Invitrogen; Cat. No. A11011) at 1:400; 488 goat anti-chicken (Invitrogen; Cat. No. A11039) at 1:400; Hoechst 33342 (Thermo Fisher Scientific; Cat. No. 62249) at 1:1000.

Slides were imaged with a Plan-Apochromat 20x/0.8 NA objective on a Zeiss LSM 880 confocal laser scanning microscope. A 7x8 (2592μm x 2954μm) tile scan with 15% overlap centered around the striatum was acquired for each section. In ImageJ (v2.3.0/1.53f), ROIs were drawn around the striatum using the DAPI and NeuN channels. NeuN and cFos channels were thresholded (500-65535 on 16-bit images). The *Analyze Particles* function (size: 30-infinity; circularity: 0.01-1.00) was run on the striatum ROI was opened on the NeuN thresholded image and the number of total particles detected was recorded. For the percentage of cFos+ neurons, the NeuN and cFos thresholded images were processed via the *Image Calculator* function with the ‘AND’ operator to isolate pixels that were double-positive for both signals in one image. The *Analyze Particles* function was used with the settings listed above to determine the number of NeuN+ cells that were also labelled with cFos. The ROIs for all the detected double-positive cells was saved and used to analyzing the cFos intensity. For cFos intensity, the ROIs from cFos+ neurons was opened on the raw 8-bit cFos image for each section. Measurements for the mean values on each ROI was recorded. Percentages and mean fluorescence intensity values were corrected by the values detected in the negative antibody control stainings.

### Ex vivo calcium imaging of striatal SPN dendrites

Mice were deeply anesthetized with a mixture of ketamine (100mg/kg) and xylazine (7 mg/kg) and perfused transcardially with ice-cold sucrose-based cutting solution containing (in mM): 184 sucrose, 25 NaHCO_3_, 1.25 NaH_2_PO_4_, 2.5 KCl, 0.5 CaCl_2_, 7 MgCl_2_, 11.6 sodium ascorbate, 3.1 sodium pyruvate, and 5 glucose (∼305 mOsm/L). Sagittal brain slices (280μm thick) were prepared using a vibratome (Leica; Model VT1200S). Slices were incubated at 34°C for 30 minutes in artificial cerebrospinal fluid (aCSF) containing (in mM): 124 NaCl, 3 KCl, 1 NaH_2_PO_4_, 2 CaCl_2_, 1 MgCl_2_, 26 NaHCO_3_, 1 sodium pyruvate and 13.89 glucose, and subsequently maintained at room temperature until recordings. All external solutions were continuously oxygenated with carbogen (95% O_2_/5% CO_2_).

Individual slices were transferred to a recording chamber and continuously superfused with aCSF without sodium pyruvate (0.5-1mL/min, room temperature). Using an Olympus BX-51-based two-photon laser scanning microscope (Ultima, Bruker), dSPNs were identified by tdTomato expression in D1-tdTomato mouse line or by the absence of eGFP in D2-eGFP mouse line, whereas iSPNs were identified by the absence of tdTomato or by eGFP expression in the dorsolateral striatum. Whole-cell patch clamp recordings were then performed from identified SPNs, aided by visualization with a 60x/0.9 NA water-dipping objective viewed through a NIR1 filter (775nm.100nm; Olympus), a Dodt contrast tube (Luigs & Nuimann), and a 2X parfocal magnification chamber (Bruker) onto a DCC3240 CMOS camera (Thor Labs) displayed with *MicroManager* software (UCSF; Vale lab). Patch pipettes (3-5 MΩ) were filled with an internal solution containing (mM): 115 K-gluconate, 20 KCl, 1.5 MgCl2, 5 HEPES, 2 Mg-ATP, 0.5 Na-GTP, 10 Na_2_-phosphocreatine (pH 7.26; 292-293 mOsm/L), supplemented with 100 μM Fluo-4 (Thermo Fisher Scientific, F14200) and 50 μM Alexa Fluor 568 hydrazide (Thermo Fisher Scientific, A10437). Recordings were made in the current-clamp configuration using a MultiClamp 700B amplifier (Axon Instrument, USA). Signals were filtered at 10 kHz and digitized at 10 kHz. Voltage protocols and data acquisition were controlled by Prairie View 5.6 (Bruker).

After whole-cell recording configuration was established, cells were allowed to equilibrate with dyes for at least 20 min before imaging. Proximal (∼50 μm from the soma) and distal (∼100 μm from the soma) dendritic structures were visualized by the red fluorescence of Alexa Fluor 568, detected with a Hamamatsu R3982 side-on photomultiplier tube (PMT, 580–620 nm) and an 810 nm excitation laser (Chameleon Ultra I, Coherent Laser Group). Calcium transients in the green channel, evoked by somatic injection of three current steps (2nA, 2ms, 50Hz), were detected using a Hamamatsu H10770PB-40 GaAsP PMT (490–560 nm, Hamamatsu Photonics, Japan). Line-scan signals were acquired for both channels across 17 pixels along dendritic segments with a spatial resolution of 0.197 μm/pixel and a dwell time of 10 μs/pixel. Signals from both channels were background-subtracted prior to analysis. Dendritic excitability (Ca^2+^ signals) was quantified as the area under the increase in green fluorescence relative to baseline, normalized to the average red fluorescence (ΔG/R). Only recordings with comparable baseline fluorescence ratios (G_0_/R_0_) before and after sTREM2 application were included in the analysis. sTREM2 (5,000 ng/ml) was superfused at 0.4ml/hr over the slice (∼300 μm from the patched cell soma) using a syringe pump and multi-barreled perfusion manifold (Cell MicroControls). Alternatively, bath perfusion of sTREM2 at 200 ng/ml was used. Data from both application methods were pooled, as no statistical differences were observed between them.

Electrophysiology and 2PLSM data were acquired using a PCI-NI6052E analog-to-digital converter card (National Instruments) and Prairie View 5.6 software (Bruker). Calcium imaging data were analyzed using custom-written Python scripts (https://doi.org/10.5281/zenodo.17380534).

### Ex vivo hippocampal field recordings

Following isoflurane anesthesia, mice were decapitated followed by brain extraction in ice-cold oxygenated (95% O_2_, 5% CO_2_) cutting solution containing (in mM): 212.5 sucrose, 10 D-glucose, 3 KCl, 1.25 Na_2_HPO_4_, 24 NaHCO_3_, 0.1 CaCl_2_, and 5 MgCl_2_, pH 7.4. Horizontal brain slices (300 µm thick) containing hippocampus were prepared using vibratome (Campden) in the cutting solution. Slices were then transferred to oxygenated artificial cerebrospinal fluid solution (ACSF) containing (in mM): 130 NaCl, 3.5 KCl, 1.25 NaH_2_PO_4_, 24 NaHCO_3_, 10 D-glucose, 2 CaCl2, and 2 MgCl2, pH 7.4. Slices were incubated for 1 hour in 32°C and at least 30 minutes in room temperature before recording. Individual slices were transferred to an immersion recording chamber, where they were submerged in oxygenated ACSF supplemented with 10 µM picrotoxin in 3 mL/min continuous perfusion at room temperature and were identified in DIC mode by ZEISS Axio Examiner upright microscope and ORCA Fusion Digital CMOS High Resolution Camera (Hamamatsu). Field Excitatory Postsynaptic Potentials (fEPSP) were recorded through a borosilicate glass capillaries (1B150F-4, World Precision Instruments) using a PC-100 vertical pipette puller (Narishige), filled with 1M NaCl solution. The recording pipette was placed in the *stratum radiatum* of the CA1 region and concentric microelectrode was placed on Schaffer collateral pathway. Evoked fEPSPs were elicited by stimulating the slice every 10 seconds using a current that elicited a 30–40% maximal response measured as the initial slope. fEPSPs were recorded in Axon Clampex 11 (Molecular Devices) using a Multiclamp 700B amplifier (Molecular Devices), a Digidata 1550B (Molecular Devices) with 10 kHz sampling rate and 2 kHz lowpass filter and analyzed using Clampfit 11.3 (Molecular Devices). Following 30 minutes of baseline recording, either 20 ng/ml BDNF and/or 200 ng/ml sTREM2 was applied for 30 minutes to induce synaptic plasticity and recording continued for additional 1 hour. For TrkB cKO experiments, virus expression was confirmed by observation of mCherry expression in the CA1. Following 10 minutes of baseline recording, sTREM2 (200 ng/ml) was applied 20 minutes prior to the 30-minute BDNF and/or sTREM2 application to examine sTREM2 effect on BDNF-induced synaptic plasticity. Average of the first 10 minutes responses were used as baselines. The fEPSPs were calculated by slope of response (mV/ms) and normalized to the baselines. Synaptic plasticity was calculated by averaging the last 5 minutes fEPSPs.

### Coimmunoprecipitation assay

Co-immunoprecipitation was used as a protein-protein interaction assay and was performed as in previous studies (*72*). Dynabeads M-280 Tosylactivated (Invitrogen; Cat. No. 14203) were resuspended in 1mL Buffer B (prepared according to manufacturer’s recommendation) and put onto the magnet. Supernatant was discarded and the beads were resuspended in 140μL of Buffer B. 20μL of the beads was aliquoted into six separate tubes, placed on the magnet and the supernatant removed. Beads were then resuspended in 6μg mouse anti-hTREM2 (R&D Systems; Cat. No. MAB1828) or 6μg mouse IgG, then brought to a final volume of 150μL with Buffer B. Tubes were incubated at 800rpm on a roller at 37°C for 18 hours. Tubes were placed on a magnet and the supernatant was discarded, the beads were resuspended in 1mL Buffer D and incubated on a roller at 37°C for 1 hour at 800rpm. The tubes were placed on the magnet, supernatant removed and washed with 1mL Buffer E twice. The supernatant was removed and beads were incubated with 250μg cell lysate for 18 hours shaking at room temperature. Then tubes were placed on the magnet and washed in 1mL Buffer E twice then washed in 1mL Dulbecco’s PBS twice and finally resuspended in 30μL Dulbecco’s PBS and stored at −80°C until used for Western.

### Protein extraction and Western blotting

Cells were washed twice with cold DPBS (Thermo Fisher Scientific; Cat. No. 14190144) and lysed in cold RIPA buffer (Thermo Fisher Scientific) supplemented with a protease inhibitor cocktail (MilliporeSigma; Cat. No. P8340) and phosphatase inhibitor cocktails (MilliporeSigma; Cat. Nos. P5726 and P0044). Lysates were incubated on ice for 10 minutes and centrifuged at 21,000 × g for 10 minutes at 4 °C. Supernatants were collected according to the manufacturer’s instructions, and protein concentrations were determined using a Pierce BCA Protein Assay Kit. Equal amounts of protein were mixed with 4× loading buffer, boiled at 98 °C for 5 minutes, and resolved on 4–12% NuPAGE Bis-Tris SDS-PAGE gels (Invitrogen) using MES SDS running buffer (Invitrogen; Cat. No. NP0002) at 130 V for approximately 1 hour. For co-immunoprecipitation experiments, 8 µL of inputs and 15 µL of co-IP beads contianing products were loaded.

Proteins were transferred to nitrocellulose membranes (Bio-Rad; Cat. No. 1620115) at 0.36 A for 1.5 hours. Membranes were washed three times for 10 minutes each in TBS containing 0.01% Triton X-100 (TBST) and blocked for 1 hour at room temperature in 5% non-fat milk in TBST. Membranes were incubated overnight at 4 °C with primary antibodies diluted in 1% milk in TBST. Primary antibodies included rabbit anti-AKT (Cell Signaling Technology; Cat. No. 4691), rabbit anti-phospho-AKT (Cell Signaling Technology; Cat. No. 2965), rabbit anti-MAPK (Cell Signaling Technology; Cat. No. 9102), rabbit anti-phospho-MAPK (Cell Signaling Technology; Cat. No. 4370), rabbit anti-phospho-TrkB (Thermo Fisher Scientific; Cat. No. MA5-14927), mouse anti-TrkB (BD Biosciences; Cat. No. 610102; 1:500), mouse anti-hTREM2 (R&D Systems; Cat. No. MAB1828; 1:500), and anti-β-actin–peroxidase (Sigma-Aldrich; Cat. No. 3854).

After primary antibody incubation, membranes were washed three times for 10 minutes each in TBST and incubated with appropriate HRP-conjugated secondary antibodies diluted in 1% milk in TBST for 1 hour at room temperature. Secondary antibodies included goat anti-mouse HRP (Jackson ImmunoResearch Laboratories; Cat. No. 115-035-146; 1:8000) and HRP-conjugated secondary antibodies from MilliporeSigma. Membranes were washed again three times in TBST and developed using enhanced chemiluminescence (ECL; Bio-Rad; Cat. No. 1705060). Signals were detected using a Bio-Rad imaging system with exposure times of 91 or 234 seconds. Band intensities were quantified using Image Lab (Bio-Rad) or FIJI (NIH) software.

### Split luciferase complementation assay

Split luciferase complementation assay was performed similar to previous studies (*73–75*). Briefly, protein-fragment fusions were created from cDNAs for TrkB and sTREM2 fused to either 5’ or 3’ hGLuc fragments. Fusion constructs were cloned into the pcDNA3 vector (Invitrogen). HEK 293T cells plated in a 96-well plate (10000 cells/well), cultured for 24 hours, then co-transfected in a 1:1 ratio with 0.05ug of each plasmid and Lipofectamine 3000 (Thermo Fisher Scientific; Cat. No. L3000015). After 48 hours, media was replaced with serum-starved DMEM without phenol red. One and a half hours after media change, coelenterazine (25μM; Gold Biotechnology; Cat. No. CZ25), the luciferase substrate, and any treatment indicated was added to each well in the plate reader and luminescence was measured (BioTek; Synergy H1 Hybrid Reader). Fusion construct pairs are as follows: N-Gluc-TrkB/empty, N-Gluc-TrkB/C-Gluc-sTREM2, N-Gluc-TrkB^ΔECD^/C-Gluc-sTREM2, and N-Gluc-TrkB/C-Gluc-TrkB. Each well was washed with dPBS then lysed with RIPA buffer for Western blot validation.

### HEK 293T–TrkB-FL Cell Culture and TrkB Signaling Pathway Analysis

HEK 293T cells stably expressing full-length TrkB (TrkB-FL) are as previously described (*76*). Cells were maintained in Dulbecco’s Modified Eagle Medium (DMEM) supplemented with 10% fetal bovine serum (FBS) and 100 U/mL penicillin–streptomycin. For signaling experiments, cells were seeded at a density of 1 × 10⁵ cells per well in 12-well plates and cultured for 48 h. Prior to stimulation with sTREM2, BDNF, or their combination, cells were serum-starved for at least 4 h in DMEM containing 0.5% FBS. Following treatment, cells were lysed, and protein samples were analyzed by Western blotting as described above. For ELISA-based detection of TrkB activation, cells were cultured under the same conditions and analyzed using the PathScan® Phospho-TrkB (panTyr) Sandwich ELISA Kit (#7108, Cell Signaling Technology) according to the manufacturer’s instructions.

**Fig. S1.**
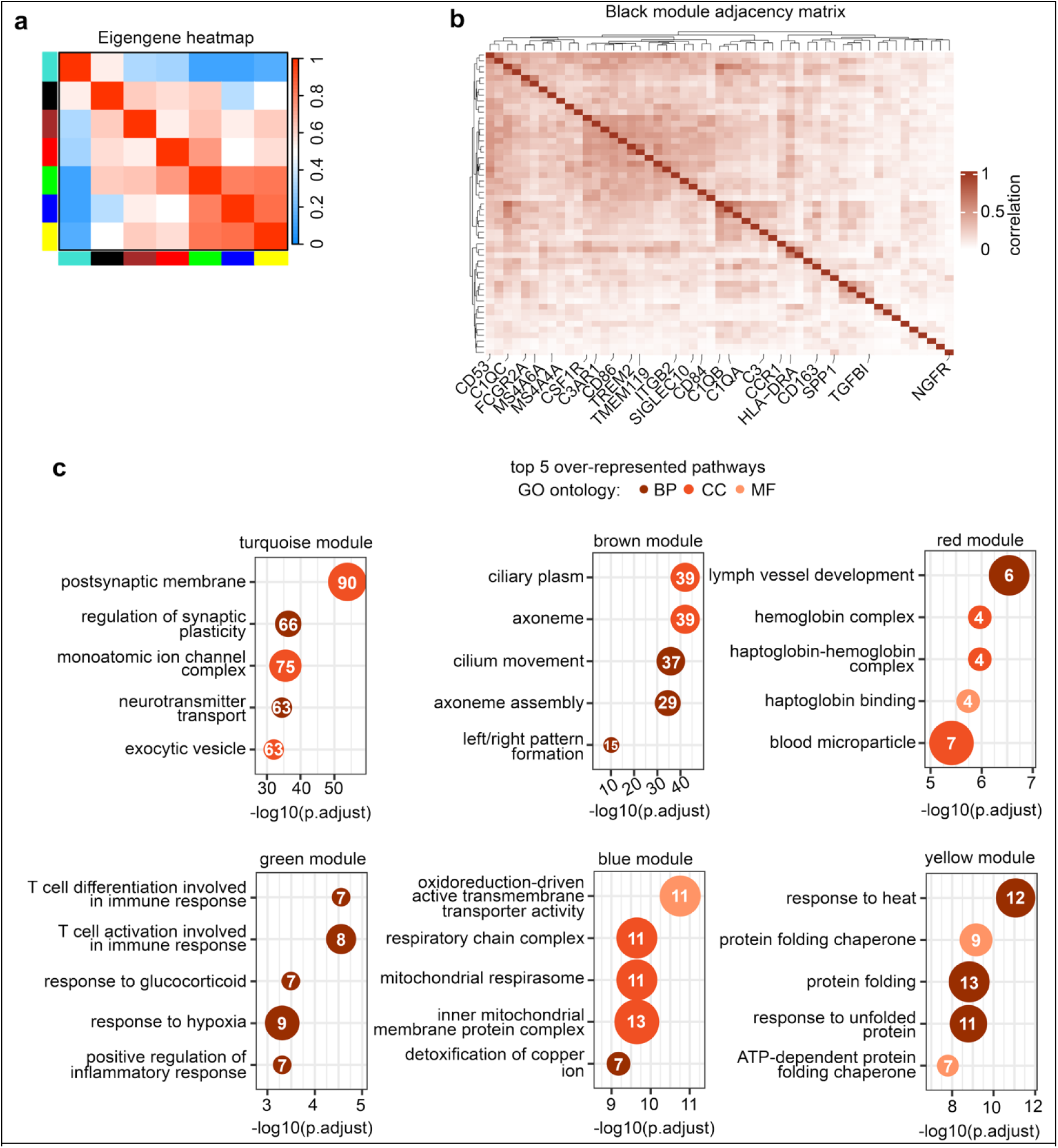
Pathway overrepresentation analysis of gene modules in human patients. (**a**) Heatmap of correlations between module eigengene values, red values on the heatmap denote higher correlation while blue values denote lower correlation. Modules are represented by color. (**b**) Adjacency heatmap of clustered genes in the black module, darker values denote higher correlation of expression between genes. (**c**) Overrepresentation analysis of GO terms using the genes assigned to each module.

**Fig. S2.**
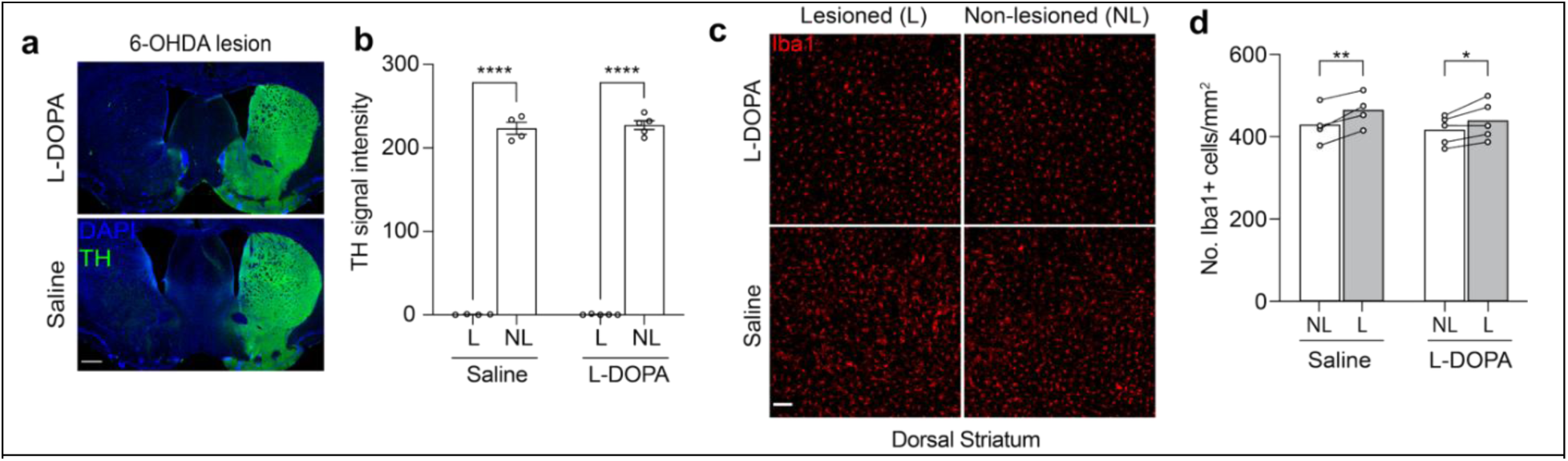
6-OHDA lesioning increases the density of Iba1+ cells in the dorsal striatum. (**a**) Representative images showing loss of TH immunofluorescence in 6-OHDA-lesioned hemispheres. Scale bar, 500μm. (**b**) Quantification (mean ± s.e.m.) of TH signal intensity in the lesion and non-lesioned hemispheres of saline- or L-DOPA-treated mice. Each dot represents the mean fluorescence intensity of 3 striatal sections per mouse (n=4-5 mice per condition). (**c**) Representative Iba1 immunofluorescence from lesioned and non-lesioned dorsal striata of saline- or L-DOPA-treated mice. Scale bar, 100μm. (**d**) Quantification (mean ±s.e.m.) of Iba1+ cell density in lesioned and non-lesioned dorsal striata. Each dot represents the average number of Iba1+ cells from 3 striatal sections per mouse (n = 4-5 mice per condition). \**p* < 0.05, \*\**p* < 0.01, \*\*\*\**p* < 0.0001, repeated measures two-way ANOVA with Sidak’s multiple comparisons test.

**Fig. S3.**
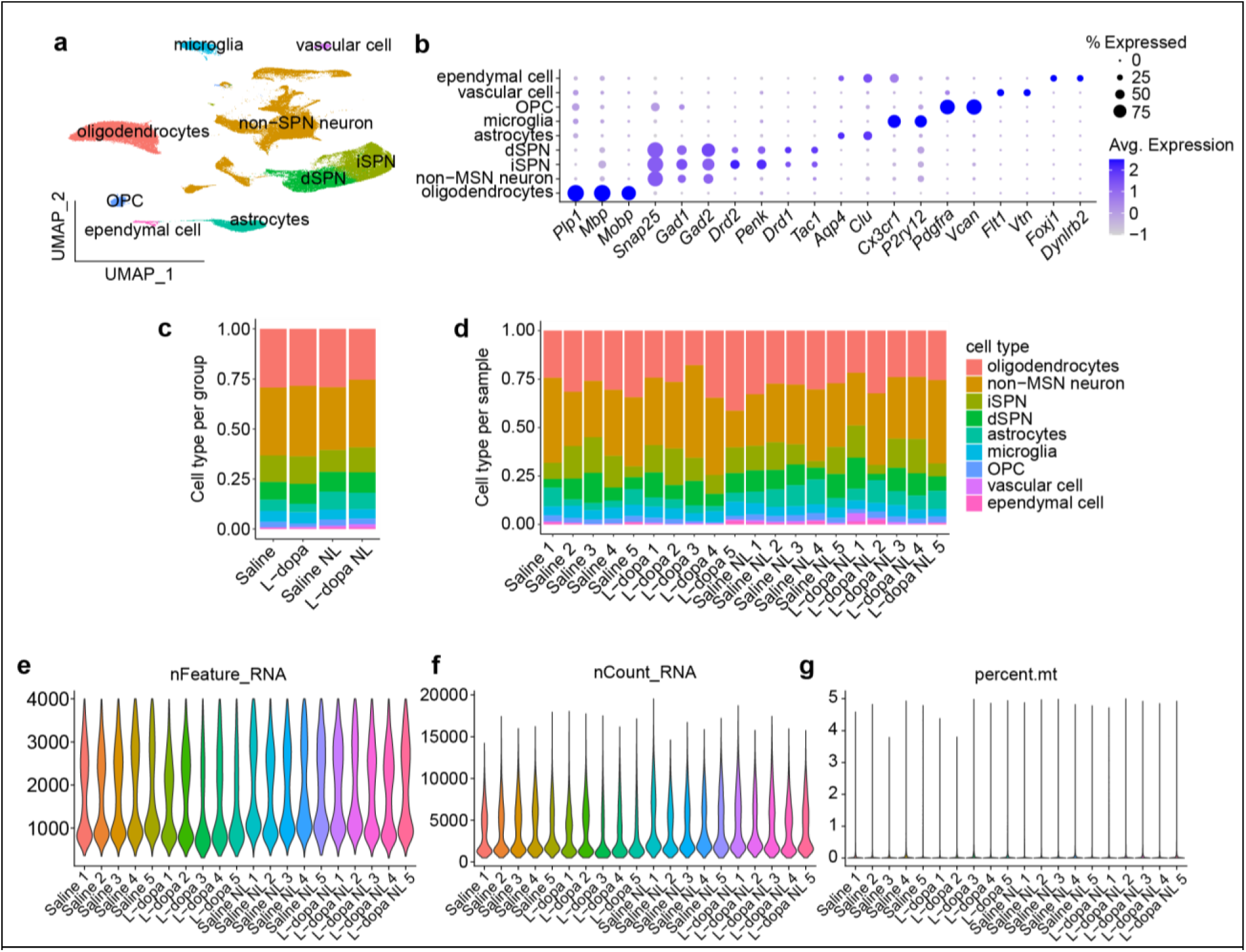
Quality assessment of snRNA-sequencing from lesioned and non-lesioned striata. **(a)** UMAP of the 191,109 nuclei sequenced from 20 striatum samples (n = 5 mice per treatment, 2 hemispheres per mouse), nuclei are colored by cell type. (**b**) Dot plot showing the percentage of (size) and average (color) expression of cell type-specific marker genes used for determining cluster identities. Stacked bar graph depicting the proportion of each cell type found within groups (**c**) and individual samples (**d**), colors correspond to allocated cell types. Violin plots showing the number of genes (**e**), counts (**f**), and percentage of mitochondrial genes (**g**) in each sample.

**Fig. S4.**
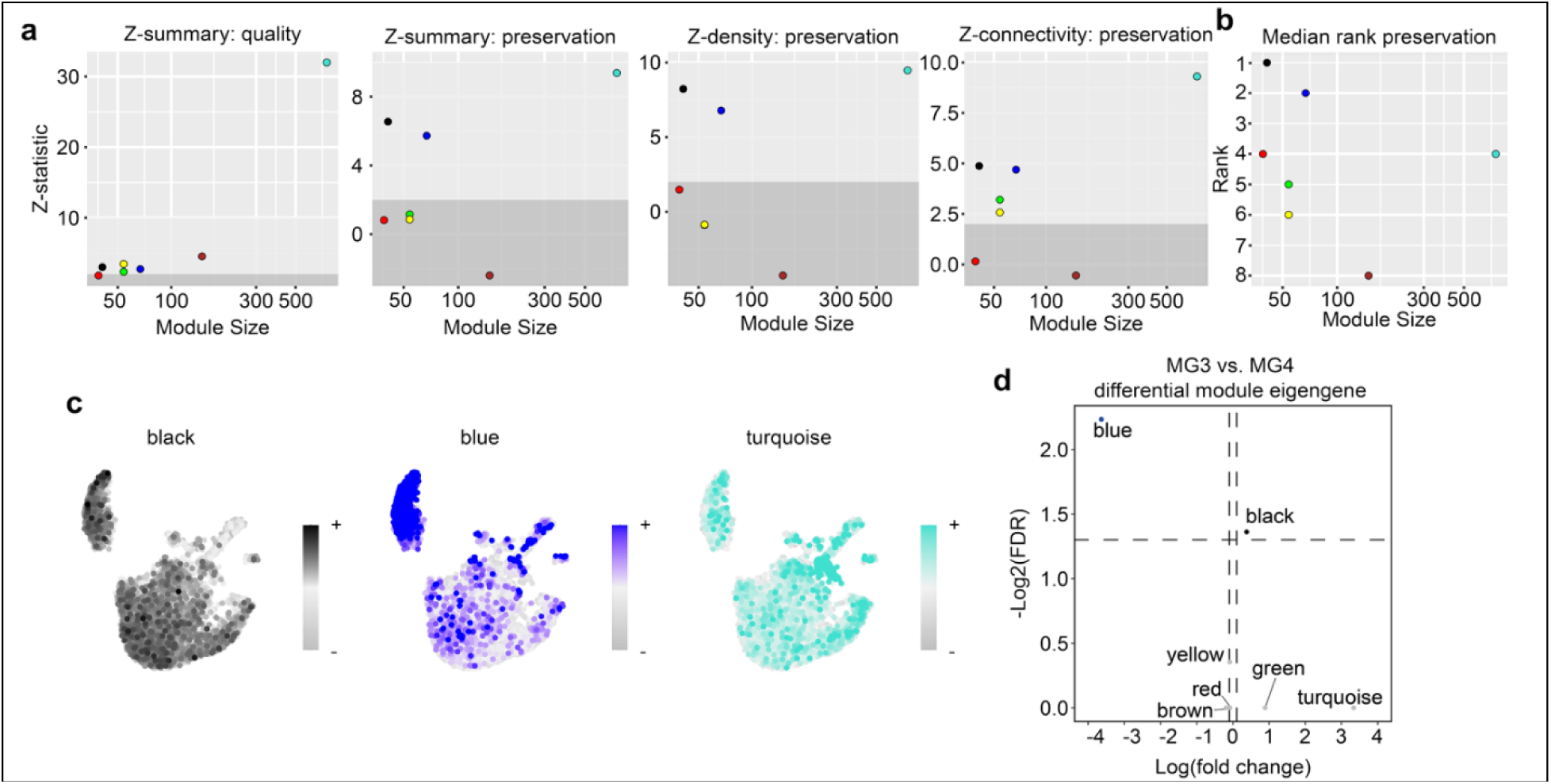
Gene networks from human PD patients are preserved in microglia from LID mice. (**a**) Scatter plots of module size versus Z-statistics calculated for module quality, preservation, density, and connectivity in mouse striatal microglia. Dot colors correspond to the respective co-expression modules. Background shades indicate thresholds for preservation strength. Z<2 are not preserved, 2<Z<10 are moderately preserved, Z>10 are strongly preserved. (**b**) Scatter plot of module size versus the median rank of observed preservation. (**c**) Feature plots of projected module eigengene values in microglia. (**d**) Volcano plot of differential module eigengenes between MG3 and MG4 subcluster. Dashed lines indicate FDR thresholds of *p*<0.05 and log_2_(fold change) thresholds of ±0.1.

**Fig. S5.**
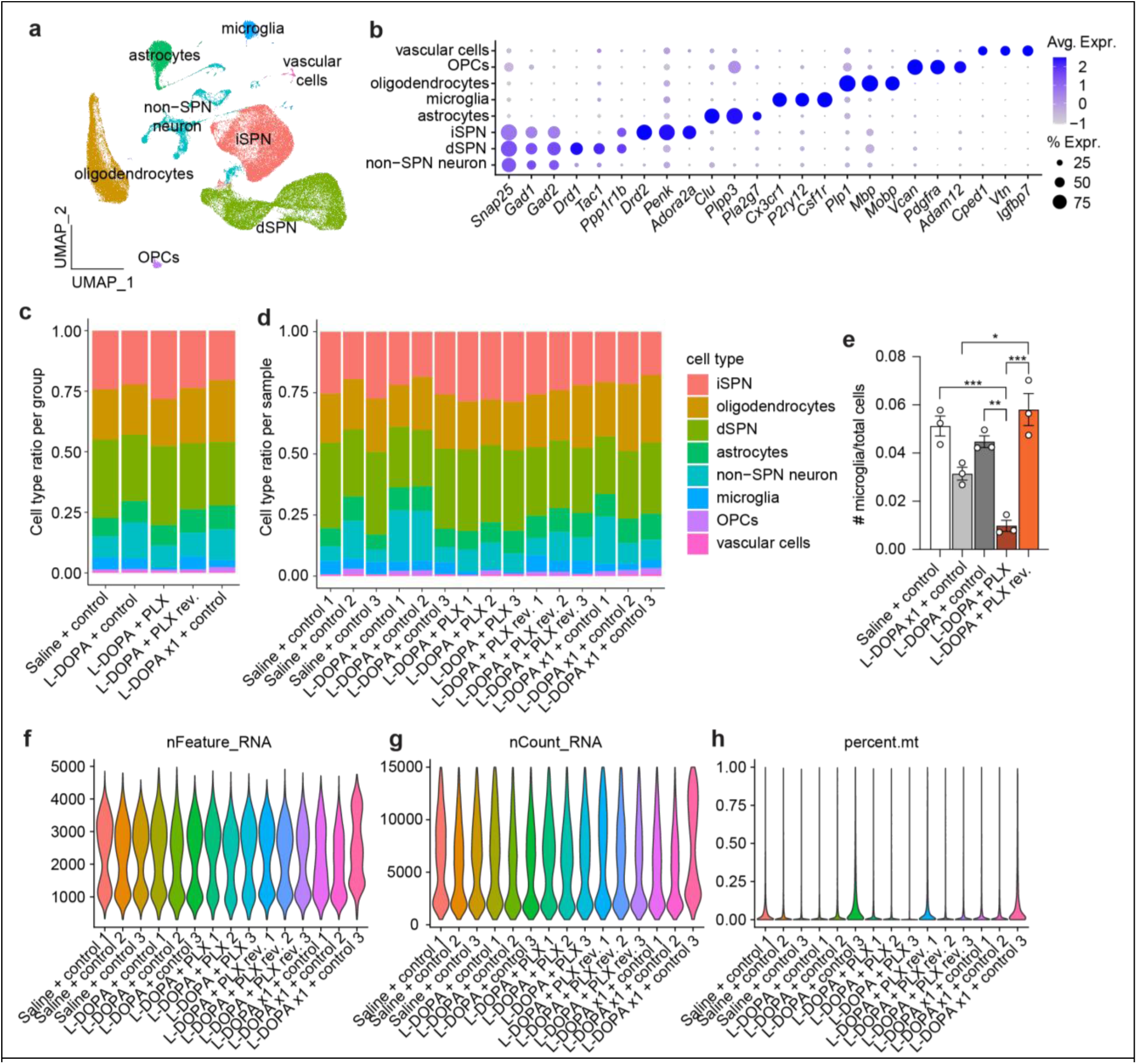
Quality assessment of snRNA-sequencing from the lesioned striatum of the microglia-depletion LID experiment. (**a**) UMAP of the 102,020 nuclei sequenced from 15 lesioned striata (n = 3 samples per group), nuclei are colored by cell type. (**b**) Dot plot showing the percentage (size) and average (color) expression of cell type-specific marker genes used for determining cluster identities. Stacked bar graph depicting the proportion of each cell type found within groups (**c**) and individual samples (**d**), colors correspond to cell types. (**e**) Bar graph showing the proportion of microglia in each group (mean ± s.e.m.), each dot represents one sample (n = 3 samples per group). \**p* < 0.05, \*\**p* < 0.01, \*\*\**p* < 0.001, 2-way ANOVA with Tukey’s post hoc multiple comparisons test. Violin plots showing the number of genes (**f**), counts (**g**), and percentage of mitochondrial genes (**h**) in each sample.

**Fig. S6.**
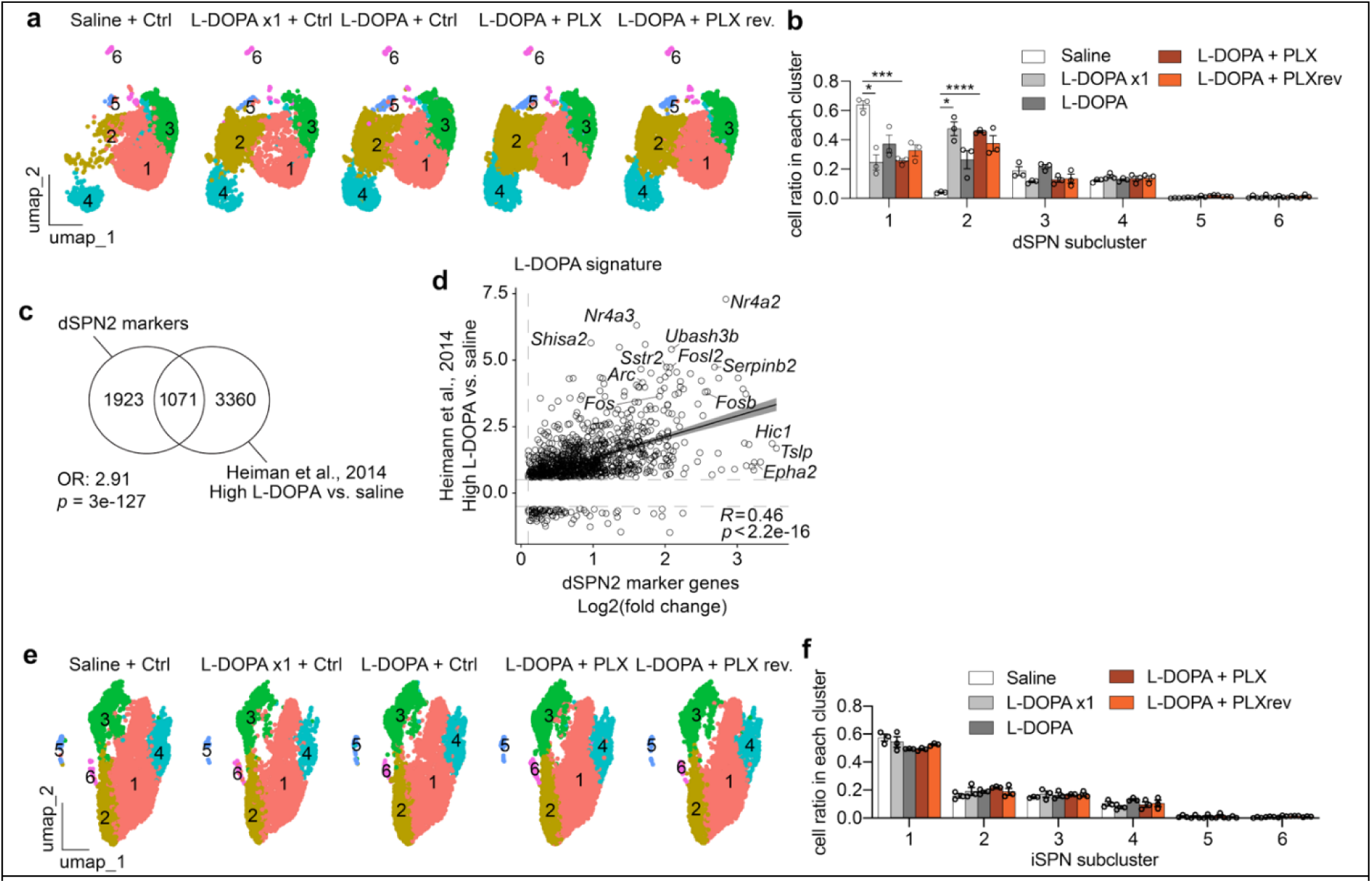
L-DOPA administration recapitulates previously published dSPN signatures. (**a**) UMAP of dSPN subclusters split by condition, color corresponds to subclusters. (**b**) Bar plot of the proportion of dSPNs assigned to each subcluster within each condition. Each dot represents the proportion of dSPNs in that subcluster in one sample (n = 3 striatal samples per group; mean ± s.e.m.). (**c**) Venn diagram of positive dSPN2 marker genes and DEGs from a previously published dataset^27^ using the same L-DOPA dosage. (**d**) Scatter plot and linear regression line (±95% confidence interval) of gene expression changes of shared DEGs from panel C. (**e**) UMAP of iSPN subclusters split by condition, color corresponds to subclusters. (**f**) Bar plot of the proportion of iSPNs assigned to each subcluster within each condition. Each dot represents the proportion of iSPNs in that subcluster in one sample (n = 3 striatal samples per group; mean ± s.e.m.).

**Fig. S7.**
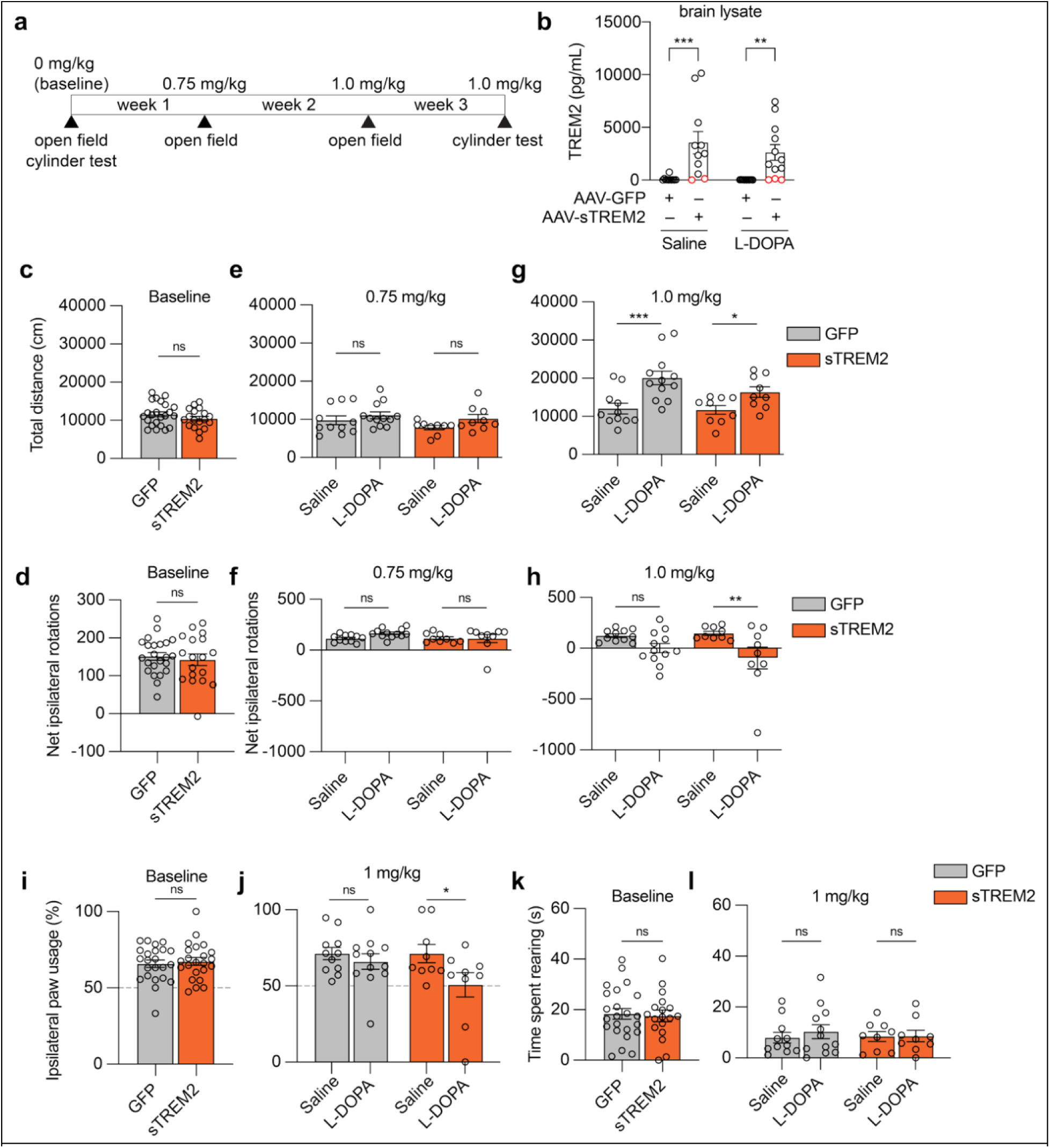
sTREM2 overexpression does not interfere with therapeutic effects of L-DOPA. **(a)** Schematic of the timeline and dosages used for behavior assays after the unilateral 6-OHDA lesioning, confirmation, and injections. (**b**) Bar graph depicting the levels of TREM2 detected via immunoassay in brain lysate samples. Each dot represents the TREM2 levels in one sample, run in duplicate. Red dots denote samples that did not reach the threshold for inclusion in the sTREM2 overexpression group (n = 5 sTREM2-injected mice). Group sizes are as follows: n = 11 GFP-injected saline-treated mice; n = 12 GFP-injected L-DOPA-treated mice; n = 11 sTREM2-injected saline-treated mice; n = 12 sTREM2-injected L-DOPA-treated mice. Bar plots of baseline measurements of total distance traveled (**c**) and net ipsilateral rotations (**d**) during open field. Total distance traveled (**e**) and net ipsilateral rotations (**f**) during open field after 0.75mg/kg administration of L-DOPA. Total distance traveled (**g**) and net ipsilateral rotations (**h**) during open field after administration of 1.0mg/kg L-DOPA. Percentage of ipsilateral paw usage during baseline (**i**) and after 1mg/kg L-DOPA (**j**) during the cylinder test. Time spent rearing at baseline (**k**) and after 1mg/kg L-DOPA (**l**) during the cylinder test. For b, e-h, j, and l, *p < 0.05, **p < 0.01, ***p < 0.001, two-way ANOVA with uncorrected Fisher’s LSD for multiple comparisons. For c, d, i, k, two-tailed unpaired t-test.

**Fig. S8.**
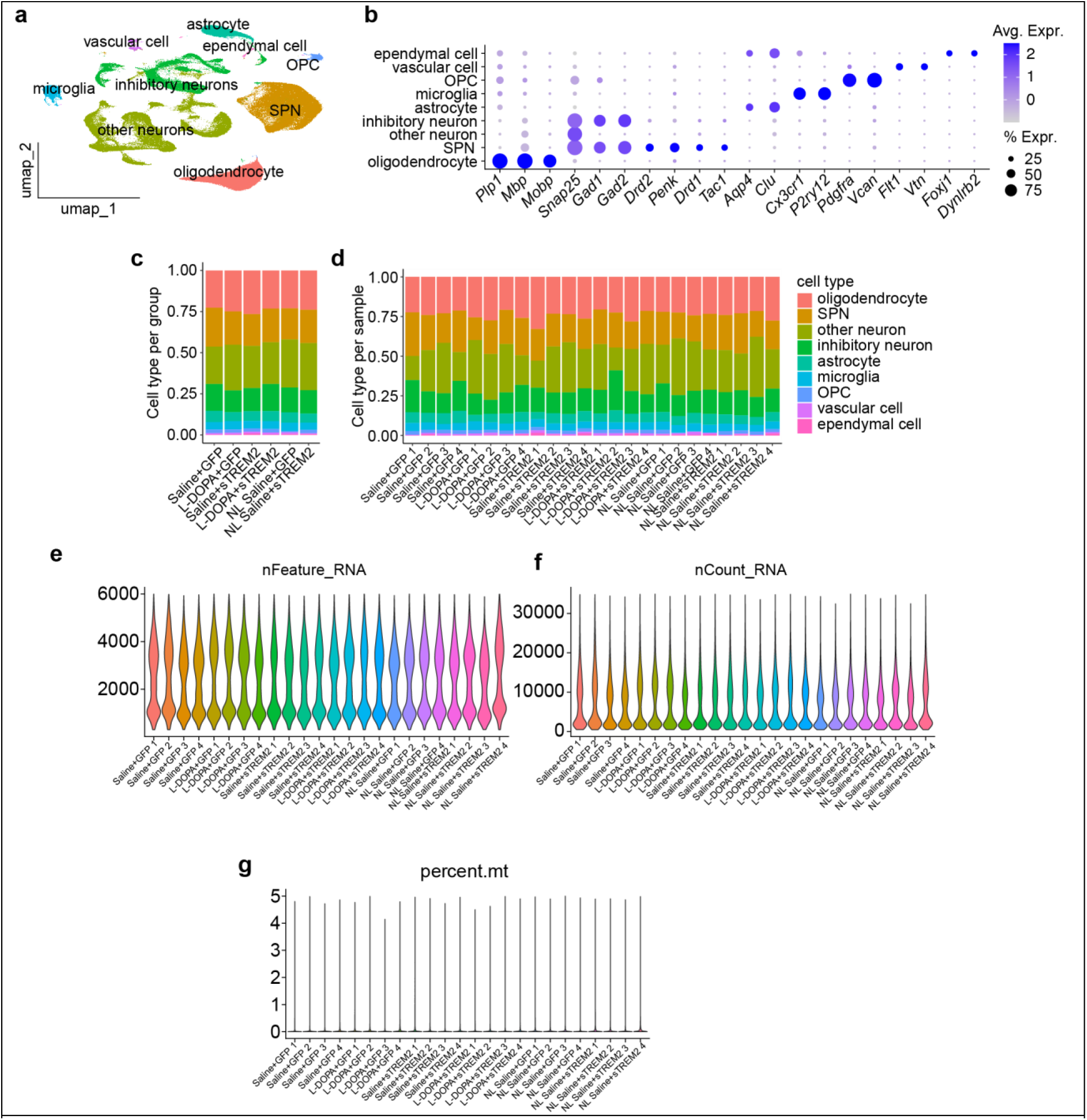
Quality assessment of snRNA-sequencing from lesioned and non-lesioned striatal samples from the sTREM2-overexpression experiment. (**a**) UMAP of nuclei sequenced from 24 lesioned and non-lesioned striata (n = 4 samples per group), nuclei are colored by cell type. (**b**) Dot plot showing the percentage (size) and average (color) expression of cell type-specific marker genes used for determining cluster identities. Stacked bar graph depicting the proportion of each cell type found within groups (**c**) and individual samples (**d**), colors correspond to cell types. Violin plots showing the number of genes (**e**), counts (**f**), and percentage of mitochondrial genes (**g**) in each sample

**Fig. S9.**
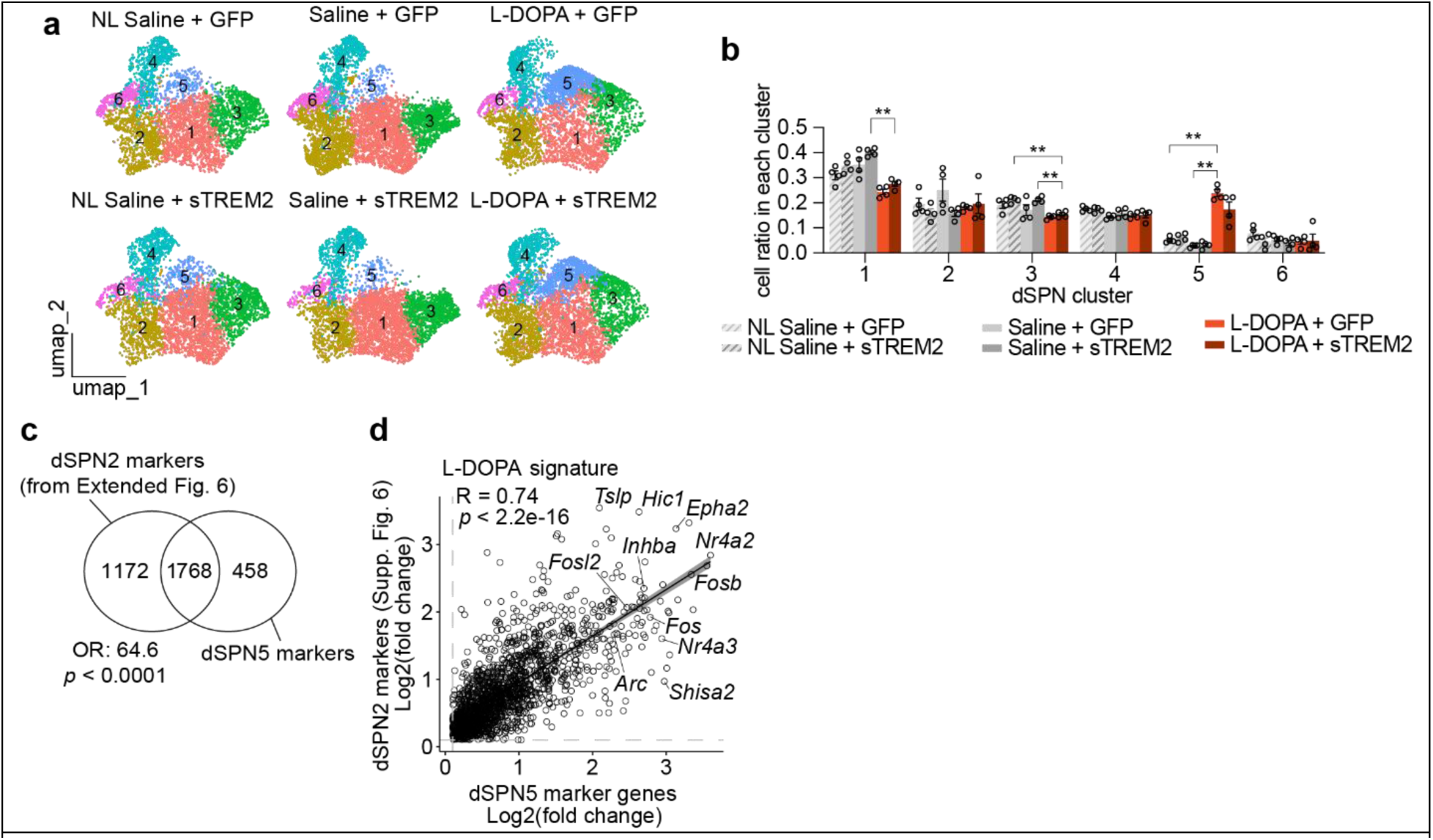
L-DOPA dSPN signatures are preserved across experiments. (**a**) UMAP of dSPN subclusters split by condition, color corresponds to subclusters. (**b**) Bar plot of the proportion of dSPNs assigned to each subcluster within each condition. Each dot represents the proportion of dSPNs in that subcluster in one sample (n = 4 striatal samples per group; mean ± s.e.m.). (**c**) Venn diagram of dSPN5 marker genes and the previous dSPN2 marker genes (from Figure 2.6). (**d**) Scatter plot and linear regression line (±95% confidence interval) of gene expression changes of shared DEGs from panel c.

**Fig. S10.**
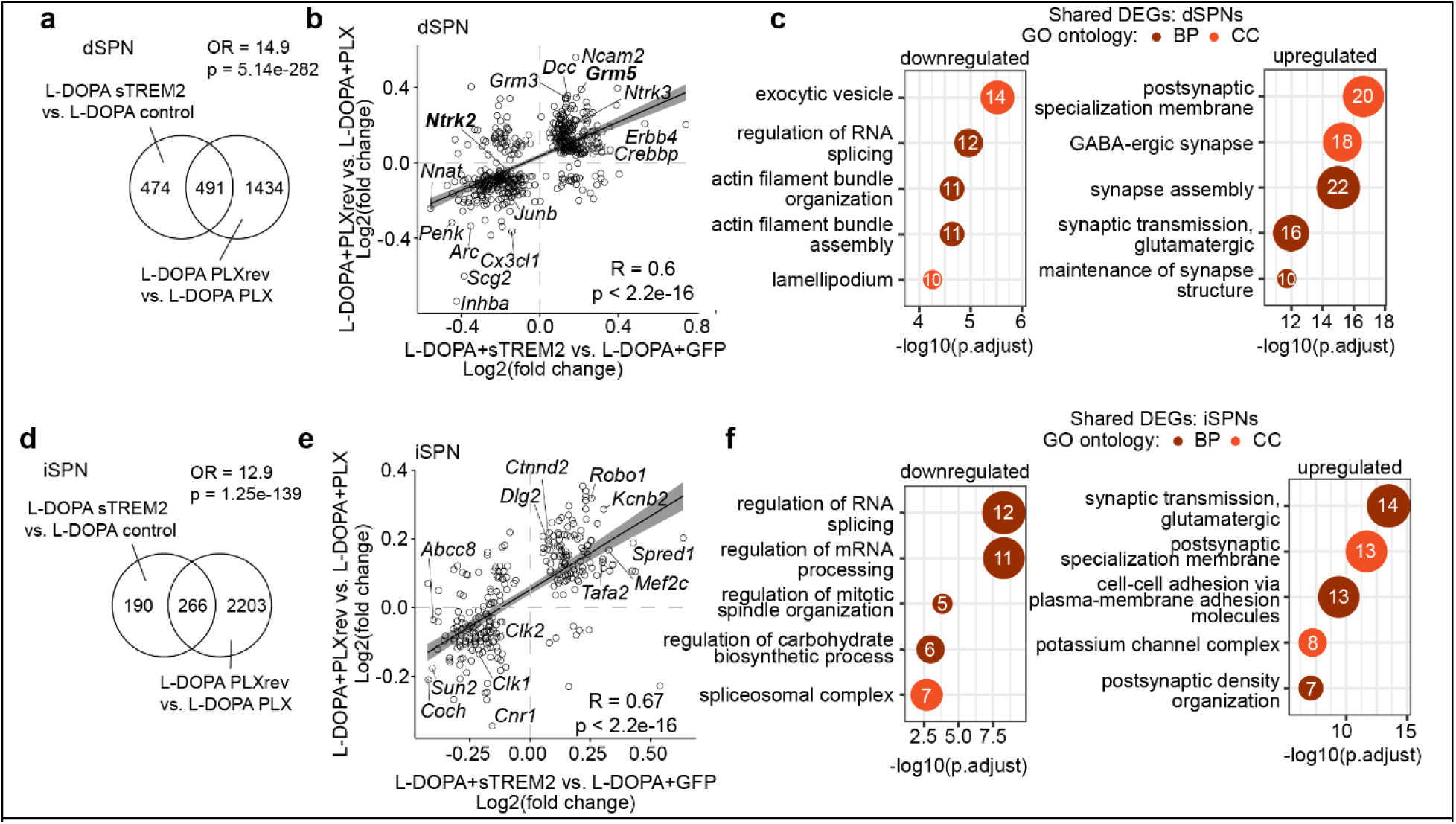
sTREM2 overexpression elicits similar SPN transcriptomic changes as microglia repopulation. (**a**) Venn diagram of pseudobulk DEGs from dSPNs in L-DOPA-treated mice injected with sTREM2 versus GFP and from L-DOPA-treated mice with microglia repopulation versus depletion. (**b**) Scatter plot and linear regression line (±95% confidence interval) of gene expression changes of shared DEGs from panel **a**. (**c**) Bubble plot of the top 5 over-represented GO terms from the up- and downregulated shared DEGs in dSPNs. Size and numbers in the bubble correspond to the number of genes in that pathway. (**d**) Venn diagram of pseudobulk DEGs from iSPNs in L-DOPA-treated mice injected with sTREM2 versus GFP and from L-DOPA-treated mice with microglia repopulation versus depletion. (**e**) Scatter plot and linear regression line (±95% confidence interval) of gene expression changes of shared DEGs from panel D. (**f**) Bubble plot of the top 5 over-represented GO terms from the up- and downregulated shared DEGs in iSPNs. Size and numbers in the bubble correspond to the number of genes in that pathway.

**Fig. S11.**
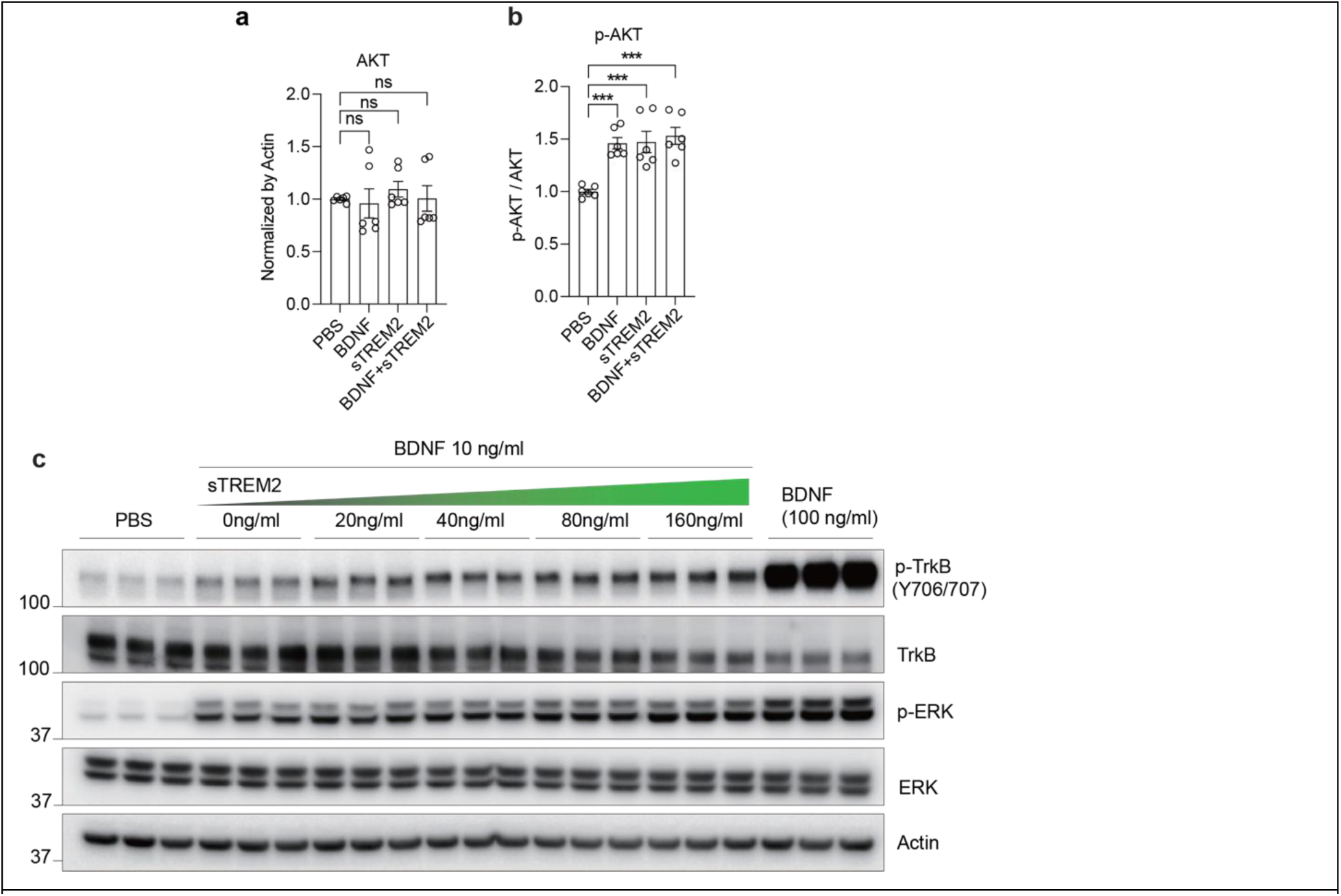
sTREM2 treatment of HEK 293-TrkB cells. Quantification of AKT (**a**) and p-AKT (**b**) relative to actin in HEK 293-TrkB cells after treatment with PBS, BDNF (20ng/mL), sTREM2 (200ng/mL), or both BDNF and sTREM2 (n = 6 samples per treatment) for 10 minutes. (**c**) Western blot of p-TrkB, TrkB, p-ERK, ERK, and Actin of lysates from HEK-TrkB cells treated with 10 ng/ml of BDNF in the presence of increasing amount of sTREM2, or 100 ng/ml of BDNF alone for 10 minutes.

### Supplementary Tables (tables in Excel format)

**Table S1. Gene expression module assignment in the basal ganglia of human PD patients**

Table containing module assignment for all of the highly expressed genes in the human bulk RNA sequencing dataset for WGCNA analysis and pathway over-representation analysis for each module.

**Table S2. Microglia subclusters from lesioned and non-lesioned LID striatal samples.**

Table containing microglia subcluster markers for each microglia subcluster and DEGs between microglia subcluster 3 and 4.

**Table S3. Gene expression in SPNs upon microglia depletion and repopulation in LID mice.**

Table containing DEGs in dSPNs and iSPNs between microglia depletion vs. control LID mice and microglia repopulation vs. depletion in LID mice and the overlapping genes between both comparisons. Also contains the dSPN subcluster markers in the microglia depletion LID experiment.

**Table S4. sTREM2-associated gene expression in LID mice.**

Table containing DEGs in dSPNs and iSPNs between sTREM2-injected vs. GFP-injected LID mice and dSPN subcluster markers for dSPN5.

**Table S5. Detailed statistics for data presented.**

Table containing detailed statistical reporting for the main figures.

